# Exact Bayesian lineage tree-based inference identifies Nanog negative autoregulation in mouse embryonic stem cells

**DOI:** 10.1101/053231

**Authors:** Justin Feigelman, Stefan Ganscha, Simon Hastreiter, Michael Schwarzfischer, Adam Filipczyk, Timm Schroeder, Fabian J. Theis, Carsten Marr, Manfred Claassen

## Abstract

The autoregulatory motif of Nanog, a heterogeneously expressed core pluripotency factor in mouse embryonic stem cells, remains debated. Although recent time-lapse microscopy data provide the unparalleled ability to monitor Nanog expression at the single-cell level, the extraction of mechanistic knowledge is precluded by the lack of inference techniques suitable for noisy, incomplete and heterogeneous data obtained from proliferating cell populations.

This work identifies Nanog’s autoregulatory motif from quantified time-lapse fluorescence lineage trees with STILT (Stochastic Inference on Lineage Trees), a novel particle-filter based algorithm for exact Bayesian parameter inference and model selection of stochastic models. We first verify STILT’s ability to accurately infer parameters and select the correct autoregulatory motif from simulated data. We then apply STILT to time-lapse microscopy movies of a fluorescent Nanog fusion protein reporter and reject the possibility of positive autoregulation. Finally, we use STILT for experimental design, perform *in silico* overexpression simulations, and experimentally validate model predictions via exogenous Nanog overexpression. We finally conclude that the protein expression dynamics and overexpression experiments strongly suggest a weak negative feedback from the protein on the DNA activation rate.

We find that a simple autoregulatory mechanism can explain the observed heterogeneous Nanog dynamics. This finding has implications on the understanding of the core pluripotency network, such as supporting the ability of mESC populations to diversify their proteomic profile to respond to a spectrum of differentiation cues. Beyond this application STILT constitutes a generally applicable fully Bayesian approach for model selection of gene regulatory models on the basis of time-lapse imaging data of proliferating cell populations. STILT is freely available at: http://www.imsb.ethz.ch/research/claassen/Software/stilt—stochastic-inference-on-lineage-trees.html

## Introduction

Nanog is a key regulator of pluripotency, whose expression is fundamentally stochastic, involving the chance synthesis, degradation and interaction of biochemical species ^1,2^. It is heterogeneously expressed ^3–5^, exhibiting strong fluctuations in expression ^4,6^ which may serve to prime mESCs for differentiation ^6,7^. Nanog binds its own enhancer as a homodimer ^8^, and Nanog-dependent feedback loops are thought to be critical to mESC regulation ^9^. However, Nanog’s mode of autoregulation has been debated. While Nanog has long been thought to exhibit positive autoregulation ^10,11^, recent studies have provided evidence for both negative feedback ^12,13^ and no direct feedback ^14^.

Nanog’s intriguing heterogeneity and its associated biological implications have motivated several deterministic and stochastic models of Nanog regulation ^15–19^. While reported deterministic models provide a population level description ^15,17^, they are unable to capture pluripotency factor heterogeneity ^20,21^ and the existence of subpopulations ^5,22–24^. Previous stochastic models describe single-cell dynamics but neglect heterogeneity arising from intrinsic noise generated by bursty production of mRNA ^25^ or slow promoter-switching dynamics ^26^ which may vary between cells. Based on the available data, a variety of mechanisms have been proposed to recapitulate Nanog’s bimodal expression distribution ^3^ including bistable switches, stochastic oscillations ^16^, and excitatory systems ^18^. To ultimately discriminate between such mechanisms, quantitative model selection based on single-cell data is required.

Time-lapse fluorescence microscopy provides a means to unambiguously label, track and quantify individual cells, thus providing critical insight into dynamics of gene expression ^27,28^. Cells may be monitored as they proliferate, thereby establishing a cellular lineage tree, capturing long-term regulatory programs such as the onset of expression of lineage-determining markers in progenitor cells ^29^ or heterogeneous response to external perturbation ^30^ at the transcript and protein level. Time-lapse fluorescence microscopy is thus well suited for investigating causative relationships between genes ^31,32^, and has recently been applied to Nanog dynamics in mESCs 4,5,14,33. However, to our knowledge, no attempt has been made to directly fit time-lapse Nanog data with stochastic dynamical models, infer parameters, and perform model selection such as of competing autoregulatory motifs.

Statistical inference based on single-cell time-lapse data presents several challenges. Stochastic models typically elude analytical solution except for simple models in the steady-state ^34,35^ or transient dynamics under simplifying assumptions ^36–38^. Simplifications of fully stochastic models, such as the linear noise approximation, are often not appropriate in the context of low copy numbers where moments are poorly estimated ^39^. Particle filter based methods for inferring the unknown copy number of chemical species and associated model parameters have been successfully applied to single-cell time series data ^40^. However, proliferating cell populations require special models capturing cellular relatedness, and observations are typically noisy and incomplete. Thus, extracting mechanistic knowledge from single-cell time-lapse fluorescence microscopy data requires methods suited to noisy, partially- and discretely-observed, heterogeneous data with small molecule numbers, small, proliferating cell populations, and intrinsic stochasticity.

To address these challenges and infer Nanog’s autoregulatory mechanism, we introduce **St**ochastic **I**nference on **L**ineage **T**rees (STILT), for fitting stochastic gene regulation models to time-lapse data of proliferating cells with known genealogy (lineage tree). STILT originally extends exact Bayesian parameter estimation and model selection for stochastic models to tree-structured data, thus enabling the investigation Nanog autoregulation using time-lapse fluorescence microscopy movies and providing a valuable, general tool for the analysis of single-cell time-lapse data. We demonstrate STILT’s capability to infer parameters and select among three models of transcriptional autoregulation: positive, negative and no transcriptional feedback. We then investigate the autoregulatory motif governing Nanog dynamics in a recently published Nanog single-cell time-lapse dataset ^4^. We compute the evidence for each model, enabling us to reject positive feedback as a likely mode of Nanog autoregulation. To resolve between no feedback and negative feedback, we design an informative perturbation experiment, predict and subsequently verify its response, finally identifying weak negative feedback as Nanog’s most probable autoregulatory motif.

## Results

### STILT: A stochastic inference algorithm using tree-structured time-lapse fluorescence microscopy data

We introduce STILT for performing parameter inference and model comparison for stochastic chemical reaction networks from fluorescence microscopy movies of proliferating cells; details are contained in the Online Methods. STILT requires as input quantitative single-cell time series data derived from time-lapse fluorescence microscopy along with the corresponding cellular lineage trees (Fig. 1A). STILT iteratively proposes new samples (particles) for both the unknown latent history of the system (including potentially unobserved species) and the distribution of parameters given the observed data. It novelly couples the bootstrap particle filter ^41^ to a model of cell division to facilitate inference of stochastic gene regulation models, compute evidences, perform model comparison, and infer parameters. By comparing proposed trajectories with the data, incompatible particles are removed, enriching the population for informative particles. Parameter posterior distributions are approximated by the particle mixture distribution after iteratively including all observations.

**Figure 1:**
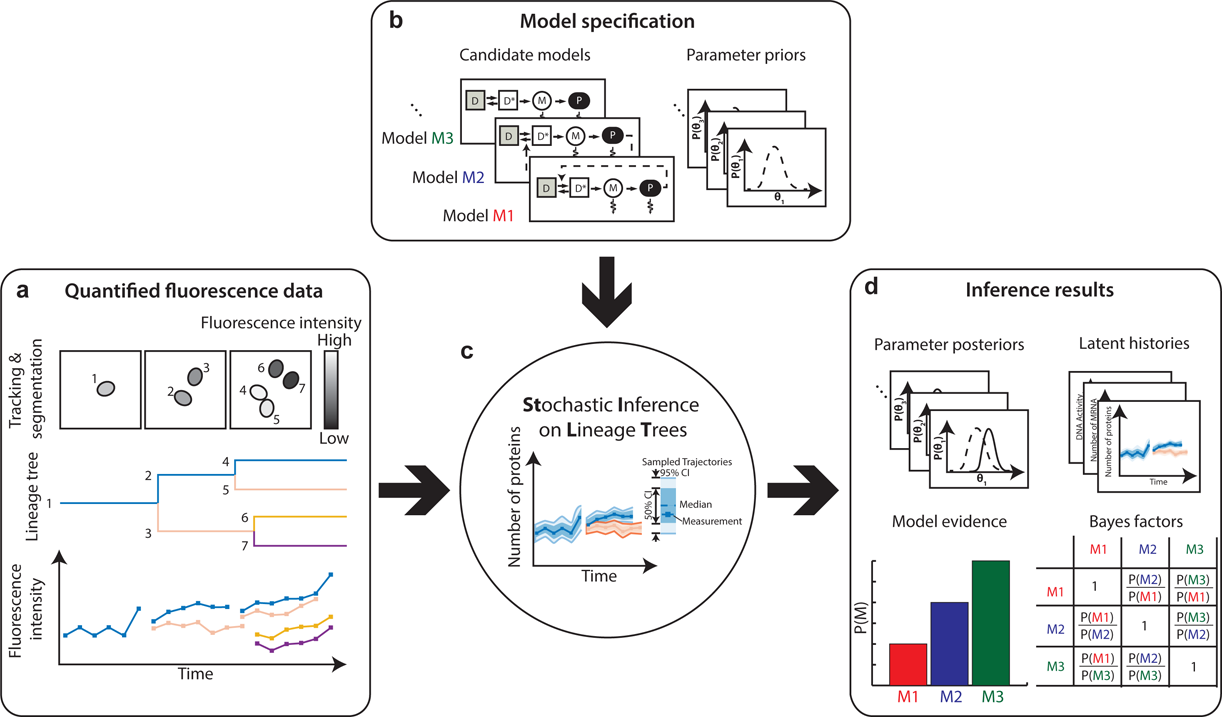
Exact Bayesian inference of stochastic gene regulation models using single-cell time-lapse fluorescence lineage trees. (A) Quantified fluorescence lineage trees are extracted from time-lapse fluorescence microscopy movies. The trees are combined with (B) candidate models and their respective parameter prior distributions and serve as input to the (C) particle filter-based inference algorithm STILT. (D) STILT generates estimates for the posterior distribution of each model parameter, latent histories, and evidence of each model. The latter is used for model comparison using Bayes Factors, which is the ratio of the marginal likelihoods of two models.

STILT requires the specification of one or more candidate models in the form of chemical reaction networks (Fig. 1B), which relate chemical species via their reactions’ stoichiometry and kinetic constants. Since the true values of parameters are generally not known, a prior distribution (i.e. from literature) is required for each parameter (Fig. 1B). STILT (Fig. 1C) then combines the experimental data, prior distributions and model structures to estimate parameters and latent histories, and approximate model evidence (Fig. 1D). Using the evidence, one can compute Bayes factors to assess the relative probability of each model and potentially reject models that cannot explain the data.

### Autoregulatory motifs

We considered three motifs for Nanog autoregulation: No Feedback (Fig. 2A), Negative Feedback (Fig. 2B), and Positive Feedback (Fig. 2C); protein affects the DNA activation/inactivation rate in case of feedback. Each model comprises a single gene in either an inactive conformation (D) with no transcription, or an active conformation (D*) with stochastic transcription, and mRNA (M) and protein (P) (see Online Methods for details). These molecular species are governed by six reactions for activation/inactivation of DNA, and production and degradation of mRNA/protein (Table 1). In the following we first validated model selection with STILT using synthetic data, and subsequently applied STILT to real Nanog time-lapse data.

**Figure 2:**
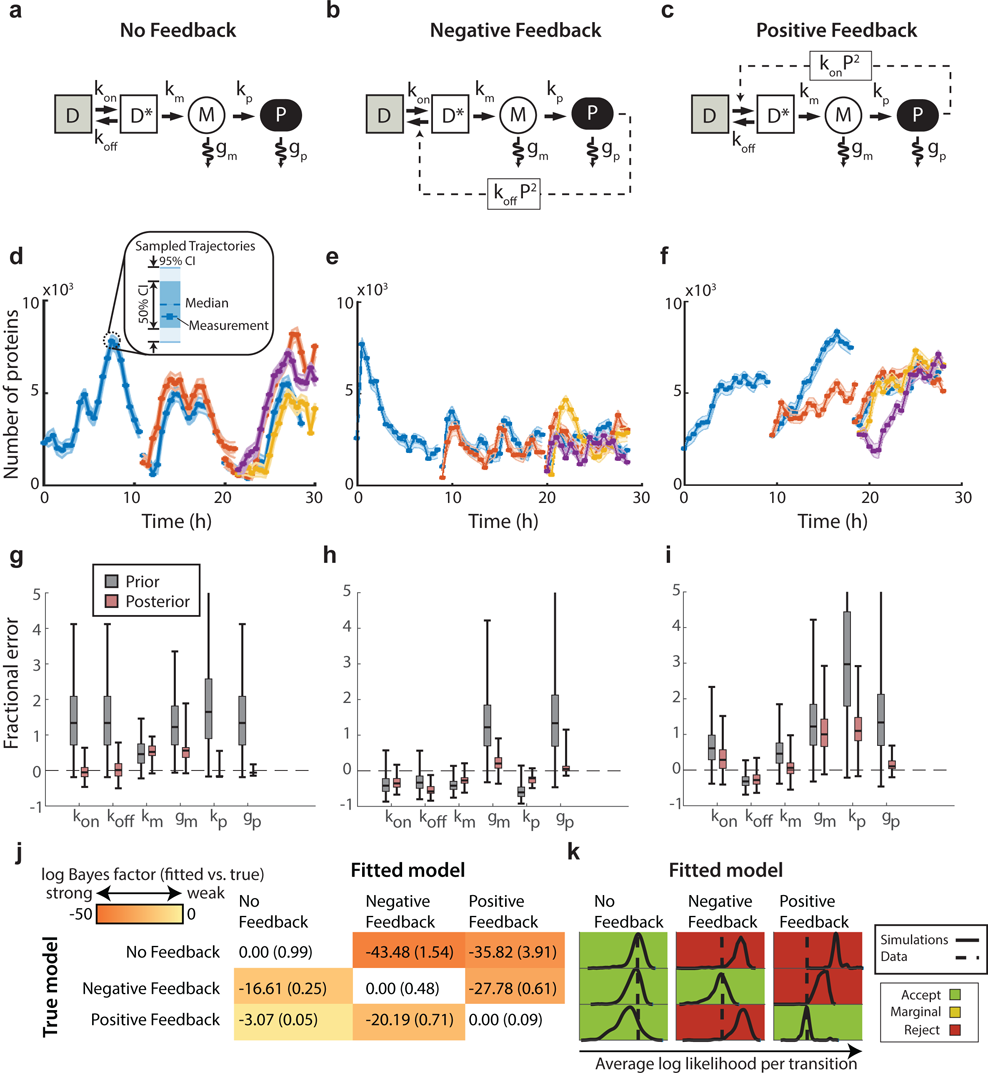
STILT correctly identifies autoregulatory models in synthetic data. We consider three simple models of transcriptional control: (A) No Feedback, (B) Negative Feedback and (C) Positive Feedback. Models differ in the propensity of DNA (D/D*) activation and inactivation. Further components of the system comprise mRNA (M) and protein (P) (see Table 1 for details on system reactions). (D-F) We simulate each model to generate quantified lineage trees of measured protein numbers, and subsequently perform inference using STILT. The median (dashed line) and 50%, 95% confidence intervals of the trajectories sampled by the particle filter (band plots) show excellent agreement with the simulated data (dots). (G-I) STILT estimates posterior distributions of model parameters (red, 99% confidence interval). For many parameters the posterior shows improved estimates compared to the prior distribution (gray) in terms of the fractional error, defined as the error of each parameter sample divided by the true value of that parameter. A fractional error of zero indicates a perfect inference result. (J) Log Bayes Factors (mean, s.d., n=3 inference runs), i.e. the difference in the marginal log likelihood P of each model from that of the true model for each dataset, indicate that the correct model is always strongly preferred (white diagonal). (K) The goodness-of-fit test (see Methods) approximates the distribution of simulated average log likelihood per transition for simulations generated using the inferred parameters for each model (solid). If the average log likelihood of the actual dataset (dashed) falls within this distribution, it indicates good agreement of the dataset with the chosen simulated model.

**Table 1:** Propensity functions for the three autoregulation models (Negative, Positive, and No Feedback) Each model is characterized by different DNA activation and inactivation reaction propensities. The remaining propensities for transcription, translation, and degradation of mRNA and protein are identical for the three models.

**Figure 4**
| Table 1 |  | Reaction Propensity |  |  |
| --- | --- | --- | --- | --- |
| Reaction | Description | No Feedback | Negative Feedback | Positive Feedback |
| DNA → DNA* | DNA activation | $k_{on} D$ | $k_{on} D$ | $k_{on} D P^2$ |
| DNA* → DNA | DNA inactivation | $k_{off} D^*$ | $k_{off} D^* P^2$ | $k_{off} D^*$ |
| DNA* → DNA* + mRNA | Transcription | $k_m D^*$ | | |
| mRNA → ∅ | Degradation of mRNA | $g_m M$ | | |
| mRNA → mRNA + Protein | Translation | $k_p M$ | | |
| Protein → ∅ | Degradation of protein | $g_p P$ | | |

### *In silico* validation

We evaluated STILT on synthetic datasets generated from the above-described autoregulatory motifs (Fig. 2A-C). We simulated each model to yield lineage trees with 3 generations and 7 cells (Fig. 2D-F, solid lines). Parameters were chosen such that cells have similar protein levels (10^3^-10^4^ molecules) in each model (Supplementary Table 1). We assume that only protein abundance was measured. Gaussian noise (=200 proteins) was added to simulate measurement error.

We applied STILT to each lineage tree using suitable priors (Supplementary Fig. 1, Supplementary Table 2), with three runs per dataset to assess robustness. Importantly, we find that STILT proposes trajectories which completely contain the observed time series for each simulated dataset (Fig. 2D-F, shaded areas); if sampled trajectories would not contain the data it would strongly indicate against that model and/or priors. The unobserved mRNA trajectories are also well inferred by the particle filter (Supplementary Fig. 2). Using fractional errors of each parameter, 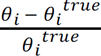, where *θ_i_^true^* is the true value of parameter *i*, we find general improvement compared to priors (Fig. 2G-I). While many parameters are estimated accurately and well contained in the posterior, some parameters are poorly identifiable, probably due to insufficient information content of the simulated data. Parameters are robustly estimated when the model is correct (Supplementary Fig. 3); inference with an incorrect model may result in local optima when the observed transitions are very unlikely to occur.

We evaluated the ability to select among different models by computing the model evidence (defined as the marginal log likelihood of the model), of each model/dataset combination. We then compute the (log) Bayes factors, i.e. differences in the evidence (Fig. 2J), and find that the true model is preferred in each case. Typically a log Bayes factor larger than 3 is considered strong evidence ^42^. For example, the difference in evidence of the (correct) No Feedback and (incorrect) Negative Feedback model (Fig. 2J, top row) is −43.48 ± 1.54 (mean ± s.d., n=3 runs), indicating strong preference for the No Feedback model. Moreover, the log Bayes factor between the correct model and the incorrect models is strong (≥3) and robust for the true model in each scenario. We compared STILT to a conventional particle filter-based algorithm which ignores cellular genealogy, inferring parameters for each cell independently. In several instances the correct model was not identified, and the Bayes factors were generally smaller when neglecting genealogy (Supplementary Fig. 4, Supplementary Table 3). Thus, STILT presents a substantial improvement over this simpler approach.

Although Bayes factors facilitate model selection, it is not in general possible to determine whether a model is “compatible” with a particular dataset, i.e., if the data could have realistically been generated by that model with the inferred parameters. Thus, we developed a simple test to assess the correctness of the inferred model and parameters by comparing the likelihood of the data with the likelihood of synthetic datasets using the assumed model and inferred parameters. We categorized each model as either reject (test statistic outside 98% confidence interval), marginal (outside 95% confidence interval), or accept (within 95% confidence interval). Our goodness-of-fit test accepts the true model for each dataset (green diagonal in Fig. 2K). By contrast, the goodness-of-fit test rejects the Negative Feedback model fit to the Positive Feedback and No Feedback datasets, and the Positive Feedback model fit to the Negative Feedback and No Feedback datasets (red), from which we can deduce that the model is unlikely to be correct for that dataset. However, the No Feedback model shows agreement with simulated datasets from both the Negative and Positive Feedback models. This level of agreement is likely due to the less constrained expression dynamics of the No Feedback model compared to the other models.

### Inference of Nanog autoregulatory motifs using time-lapse fluorescence genealogies rejects positive autoregulation

We next applied STILT to study the debated autoregulation mechanism of Nanog, using time-lapse data from a recent single-cell study ^4^. In these experiments, the fluorescence intensity of NanogVENUS, a reporter for the protein expression of the pluripotency factor Nanog, was quantified for single cells over several generations (Fig 3A). We converted fluorescence intensities in 15 subtrees (7 cells per subtree) to absolute protein numbers (see Online Methods) and performed minimal data cleaning to remove incorrectly segmented or quantified measurements (Supplementary Fig. 5). Finally we used STILT to perform inference with the three autoregulatory motifs introduced above. Prior distributions for each model parameter were estimated from available knowledge (Supplementary Table 4).

**Figure 3:**
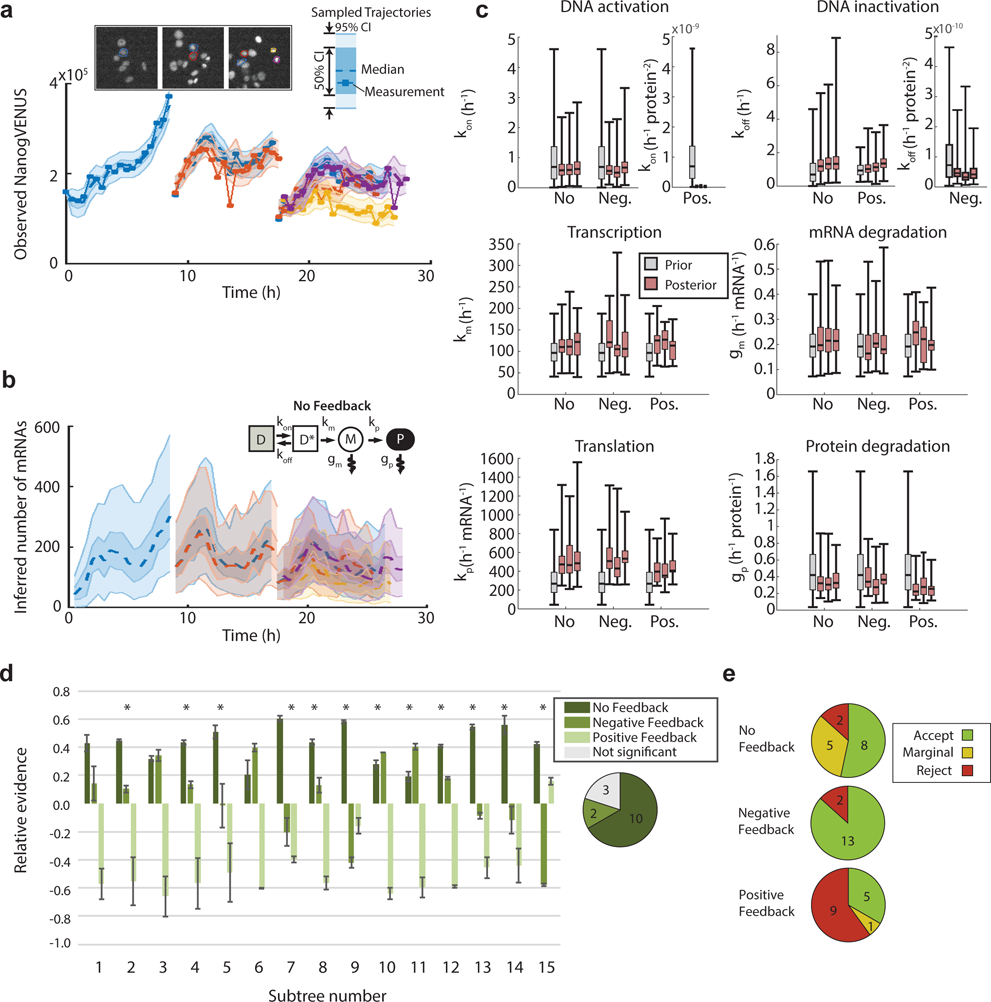
Model comparison suggests that NanogVENUS expression dynamics in ESCs are best explained by the No Feedback or Negative Feedback model. (A) STILT yields samples for the latent trajectories of proteins that reproduce and contain the observed data (shown for the No Feedback model). Colony images are shown at the first time point of each generation. (B) The latent history of mRNA is inferred and agrees with previous estimates of the mRNA copy number of Nanog in ESCs (shown for the No Feedback model). (C) We compare the estimated posterior distributions of model parameters for the same subtree fit with each of the models. We find that estimates for mRNA and protein parameters are robust between technical replicates and across models. (D) We compute the evidence of each model, shown relative to the average over all models for that subtree (mean, s.e.m., n=3), scaled by evidence range for that subtree (see Table S6 for absolute values). We find that the No Feedback model provides the largest evidence in most cases (*, significant with p<0.01), while the Negative Feedback model is preferred for four subtrees. The evidence for the Positive Feedback model is generally lower than the other models. The frequency for which each model is significantly more likely than remaining models is shown with a pie chart. (E) The goodness-of-fit test indicates that the Negative Feedback model is accepted for most subtrees (13/15) compared to 8/15 for No Feedback and 5/15 for Positive Feedback. Each model is rated as accept, marginal or reject based on the result of the goodness-of-fit test.

First, we evaluated the inference results on the Nanog time-lapse data. STILT produced sampled trajectories that agree well with the measured time series (Fig. 3A-B shows one subtree fit with the No Feedback model; see Supplementary Fig. 7 for all models and subtrees), indicating that all models are capable of reproducing the observations with the assumed parameter distributions, albeit with varying likelihoods. The estimated latent mRNA abundances (Fig. 3B) agree well with recent estimates of approximately 100-300 copies per cell ^5,43^. We find that the subtrees are informative in the sense that they cause shifts in the parameter posterior distributions relative to the priors (Fig. 3C, Supplementary Fig. 8, Supplementary Table 5). Moreover, parameters are robustly estimated over three technical replicates (Supplementary Fig. 8). Next, we estimated the evidence of each model for each subtree (Supplementary Table 6). We find that the No Feedback model is preferred in most cases (11/15), and is significantly greater than the next best model in 10 of these instances (Fig. 3D). For four subtrees the Negative Feedback model is preferred, and is significantly greater in two of these. By contrast, the evidence was consistently much lower for the Positive Feedback model.

Finally, we used the goodness-of-fit test to assess the ability of each model to explain the data. We found that both the No Feedback and Negative Feedback models agree well with the observed datasets when using the median of the estimated posteriors (Fig. 3E; Supplementary Fig. 9, Supplementary Table 7). The Negative Feedback model is compatible with the most subtrees (13/15 accepted); in contrast only 8/15 subtrees were compatible with the No Feedback model (5 subtrees were marginally accepted). However, the Positive Feedback model is accepted for only 5/15 subtrees, and marginally for one additional subtree. For two subtrees no model could be rejected, and for subtree 14 all models are rejected (Supplementary Table 7).

### Model-based experimental design for selection of Nanog autoregulation motif

Based on the goodness-of-fit test and Bayes factors analysis, we can reject positive feedback as a putative motif for Nanog autoregulation for the analyzed datasets. To discriminate between the remaining two alternatives, we used STILT to devise an experiment whose outcome would differ significantly for the No/Negative feedback models. We consider exogenous transgenic Nanog, which would increase the effective rate of DNA inactivation in the Negative Feedback model (Fig. 4A). We simulated negative feedback using the previously inferred parameters, while introducing varying levels of exogenous Nanog (P_ex_). We found a strong shift in endogenous Nanog dynamics at only a few hundred thousand molecules of exogenous Nanog, and complete down-regulation for P_ex_ > 10^6^ (Fig. 4B). By introducing exogenous Nanog we expect rapid decrease in endogenous Nanog levels for the Negative Feedback model, in contrast to constant levels for the No Feedback model (evaluated at 46h, Fig. 4C).

**Figure 4:**
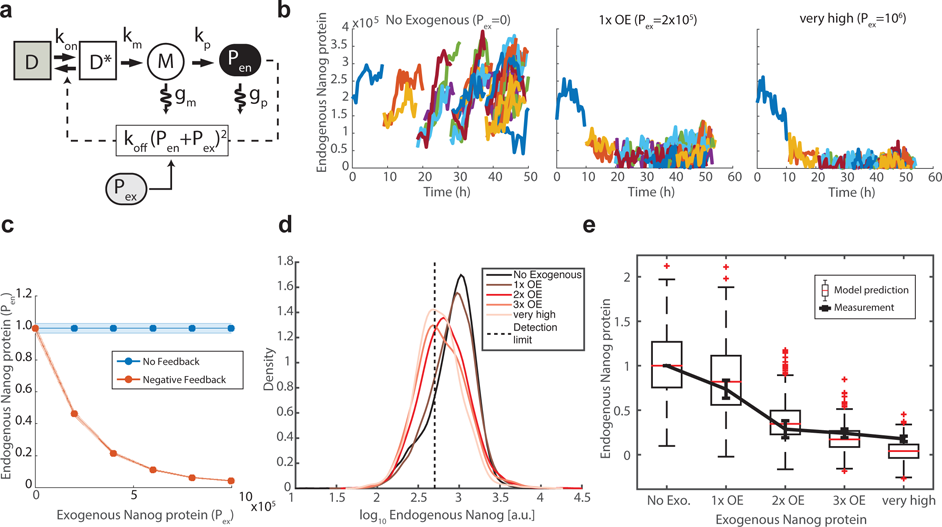
We experimentally verify the predicted response of the Negative Feedback model to Nanog overexpression. (A) We modify the Negative Feedback model to incorporate exogenous Nanog (P_ex_) which acts with endogenous Nanog (Pen) to increase the propensity of DNA inactivation. (B) Using the previously inferred model parameters we generate synthetic trees for various levels of exogenous Nanog, illustrated for 0 molecules, 2x10^5^ molecules (100% increase) and 10^6^ molecules (500% increase). (C) We predict strong downregulation (fold-change relative to median expression of unperturbed cells) of endogenous Nanog (mean, ± 2 s.e.m., n=30 simulations) for the Negative Feedback model; the No Feedback model is unperturbed by exogenous Nanog. (D) Endogenous Nanog levels decrease as the amount of exogenous Nanog increases. Detection threshold shown as dashed line (see Figure S10A). (E) Using the Negative Feedback model with exogenous Nanog, we compare the predicted fold-change (box-and-whiskers) in endogenous Nanog in response to exogenous Nanog overexpression with the experimentally determined median fold-change (mean, s.e.m. of 3 replicates) (line). Fold-change is relative to median expression level of endogenous Nanog in the unperturbed (No Exogenous) compartment.

We tested our model prediction using an mESC line with fluorescent reporters for both endogenous and exogenous Nanog (Supplementary Fig. 10A). We quantified exogenous Nanog and defined 5 compartments of expression: No Exogenous, 1x, 2x, and 3x overexpression (OE), and “very high”, relative to endogenous expression in untransfected cells (Supplementary Fig. 10B,C). In agreement with recent reports ^12,13^, expression of transgenic exogenous Nanog was found to induce a dose-dependent down-regulation of endogenous Nanog production (Fig. 4D). We then replicated the experimental perturbation using STILT. Using the estimated parameters, we simulated exogenous Nanog corresponding to the quantity of exogenous Nanog of each overexpression compartment. We found excellent agreement between the predicted and measured decrease in endogenous Nanog expression levels upon perturbation (Fig. 4E). Note that the prediction uses only parameters inferred from the time-lapse data and the estimated quantities of exogenous Nanog. The agreement suggests that the data are well explained by negative feedback with the assumed mechanistic model, and not by the No Feedback model.

### Comparative validation of estimated parameters for the Negative Feedback model

STILT yields estimates of parameters including the rate of switching between active and inactive DNA conformations, transcription and translation rates, and degradation rates of mRNA and protein. These estimates agree well with previous estimates. The inferred median mRNA degradation rate for each subtree ranges from 0.1145-0.4255 (mean 0.2262, n=15) per molecule per hour, which agrees well with the previous estimate of 0.147 ^14^ (see Online Methods). Protein degradation rates range from 0.0310-0.4410 (mean 0.2197) per molecule per hour, consistent with the previous estimates of 0.14-0.35 ^4,44^. Thus, both Nanog protein and mRNA have a comparable half-life of ~3 hours. Transcription rates range from 67.2-181.8 (mean 110.3) per hour, consistent with the estimate of 126.6 per hour ^14^. Translation rates range from 215.9-1142.0 per mRNA per hour. This quantity is not well characterized in literature, but agrees roughly with the estimate of up to 1000 estimated for mouse fibroblasts ^45,46^. The mean value of these estimates across subtrees is similar between the No Feedback and Negative Feedback models: 115.0 vs 110.3 for translation; 0.258 vs 0.226 for mRNA degradation; 637.8 vs 619.5 for translation; and 0.241 vs 0.220 for protein degradation, for the No Feedback and Negative models, respectively.

DNA activation and inactivation rates cannot be easily assessed since they represent an abstraction of a more complicated biochemical process. For example, activation might correspond to changes in the DNA and histone modification state of the promoter which permit greater transcriptional activity ^5^. Nonetheless, the estimated rate of activation ranges from 0.2737-1.737 (mean 0.6854) per hour, which is consistent with the estimate of 1.692 per hour in the simple unregulated telegraph model of Ochiai *et al.* ^14^. The inactivation rate ranges from 0.1338x10^−11-^1.123x10^−10^ (mean 5.6910x10^−11^) per hour. In the Negative Feedback model this rate scales quadratically with the number of proteins to give a total rate of approximately 2.5-5.0 per hour (assuming 2-4x10^5^ Nanog protein molecules per cell). This estimate is substantially smaller than the estimate of 36.54 per hour in the telegraph model ^14^. However, there the model assumes DNA to be inactive whenever active transcription is not detected. In contrast, the stochastic nature of our model allows DNA to remain in the active state even between transcription events, which may contribute to a reduced overall rate of DNA inactivation. We also note that the estimated number of mRNAs inferred by STILT, which ranges from approximately 0-300, agrees well with previous estimates of approximately approximately 100 ± 100 ^5,43,47^, see Supplementary Fig. 7. In summary, STILT achieves comprehensive rate constant estimates of the different processes governing Nanog dynamics solely from a time-lapse study. These are in good agreement with the results from various dedicated studies, each independently focusing on selected aspects such as DNA (in-) activation or mRNA/protein synthesis and degradation.

## Discussion

A variety of hypothetical mechanisms for Nanog regulation have previously been proposed, including bistability or oscillations ^16^, and excitatory excursions from a stable state ^18^. Such mechanisms can produce heterogeneous steady state distributions similar to those observed in snapshot experiments ^3,48^. On the other hand, Nanog transcriptional dynamics have been described statistically using a simple unregulated telegraph model, fit to the timing of periods of gene activity ^14^. However, until now extracting mechanistic knowledge from fluorescent fusion protein trajectories has been hampered by the lack of suitable inference techniques. In particular, the intrinsic stochasticity of Nanog expression at the single-cell level and the proliferating nature of mESC populations necessitate an approach that is fully stochastic, Bayesian, and suited to tree-structured data. Using STILT, we overcome these challenges to make use of the full information content of time-lapse fluorescence movies, and quantitatively fit and select among putative models of autoregulation.

Interestingly, STILT indicates greater evidence for the No Feedback model for many subtrees, and Negative Feedback for fewer subtrees; Positive Feedback consistently has the lowest evidence. However, the goodness-of-fit test indicates superior agreement with data for the Negative Feedback model. The stronger evidence for No Feedback arises because the fitted parameter values are *a priori* more likely with the assumed priors compared to those of the Negative Feedback model; the Negative Feedback model agrees with the data for a more limited set of parameters, which were assumed to be less likely. However, as for all Bayesian inference methods, this result is influenced by the choice of priors and thus should be considered in context of the goodness-of-fit test results.

To discriminate between No Feedback and Negative Feedback, we used STILT as an experimental design tool, and quantitatively predicted the strength of down-regulation upon overexpression. Further investigation using novel experiments revealed the expected strong down-regulation upon high expression of transgenic Nanog, in very good agreement with model predictions. Taken together, we conclude that Nanog negative autoregulation is indeed likely, but has a prominent effect only at relatively high levels of protein expression, which renders model discrimination based on Bayes factors alone difficult. The lack of strong autoregulation suggests stable oscillations ^16^ to be unlikely, in accordance with previous analysis ^4^, and supports the notion that Nanog undergoes broad fluctuations which serve to diversify the mESC population’s ability to respond to differentiation cues ^6,7^.

The inferred motif naturally represents a simplification of Nanog’s true regulatory mechanism. For example, although Nanog autoinhibition is thought to be mediated by Zfp281 and the NuRD complex ^12^, these factors are omitted for simplicity; this is equivalent to assuming Zfp281 abundance to be approximately constant. We further neglect the possibility of monoallelic expression of Nanog. However, it has been previously shown that Nanog that both Nanog mRNA and protein are highly correlated between alleles ^47,48^, motivating this assumption. Despite these simplifications, the Negative Feedback model i) produces sample trajectories which reproduce the observed data, ii) agrees quantitatively with observed fluorescence lineage trees using the goodness-of-fit test, and iii) accurately predicts the magnitude of downregulation in overexpression experiments. Thus we conclude that the autoinhibitory motif provides a simple but accurate description of Nanog protein dynamics, superior to the considered alternatives.

Fitting mechanistic models to time-lapse data facilitates the analysis of latent variables and enables the design of informative experiments. The sampled trajectories provide valuable insight into the dynamics of latent variables, including DNA activity and mRNA copy number. The inferred trajectories can also be analyzed to provide information about gene activity, such as inferring continuous versus bursty transcription, possible oscillations, refractory periods, etc. ^49^ For example, examining the mRNA trajectories (Supplementary Fig. 7) we observe both burst-like and sustained transcriptional modes.

Lastly, while we have focused on Nanog autoregulation, STILT may be used for inference and model selection for arbitrary stochastic gene regulation models applied to fluorescence lineage trees (e.g. in *B. subtilis* or *E. coli* ^31,50^), thus enabling quantitative and exact analysis of lineage-tracked time-lapse fluorescence data. The generic MATLAB implementation is provided as open source with SBML compatibility for easy import of user-specified models.

## Author contributions

TS and FJT conceived the interdisciplinary approach. CM and MC designed and supervised the study. JF designed, developed and implemented STILT. SG developed the STILT package. JF performed simulations and analysis. TS, SH planned and SH performed the exogenous Nanog experiments. FJT and TS provided critical comments. JF, CM, MC wrote the manuscript.

## Acknowledgments

We thank Virginia-Cezara Luca for single cell quantification, Will MacNair for feedback on the manuscript and analysis suggestions, and Jan Hasenauer & Michi Strasser for helpful comments and insightful discussions. We also thank the Helmholtz Zentrum München, ETH Zürich, and the ERC (starting grant Latent-Causes to FJT) and the SystemsX.ch (RTD HDL-X) for generous funding support.

## Supplementary Figure captions

**Figure S1.**
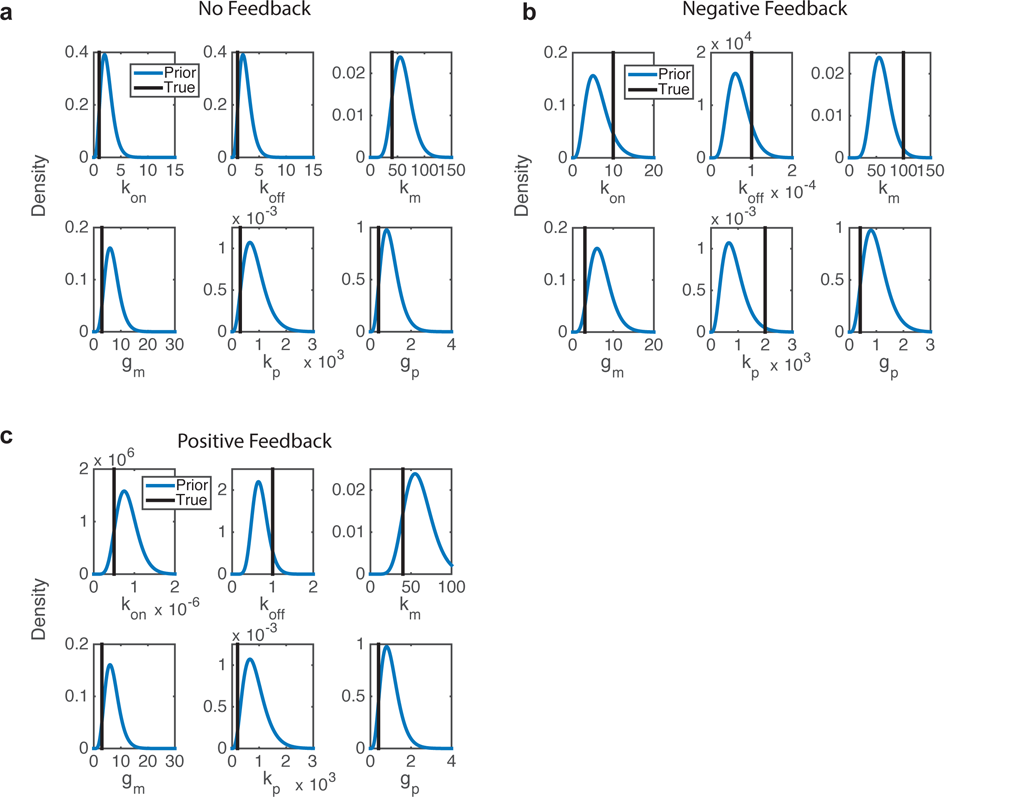
Prior distributions for *in silico* testing. STILT was run using parameter prior distributions tailored to each model, chosen such that the true model parameter used for data generation is contained in the distribution, but not identical to the mode of the distribution. (A) No Feedback, (B) Negative Feedback and (C) Positive Feedback.

**Figure S2:**
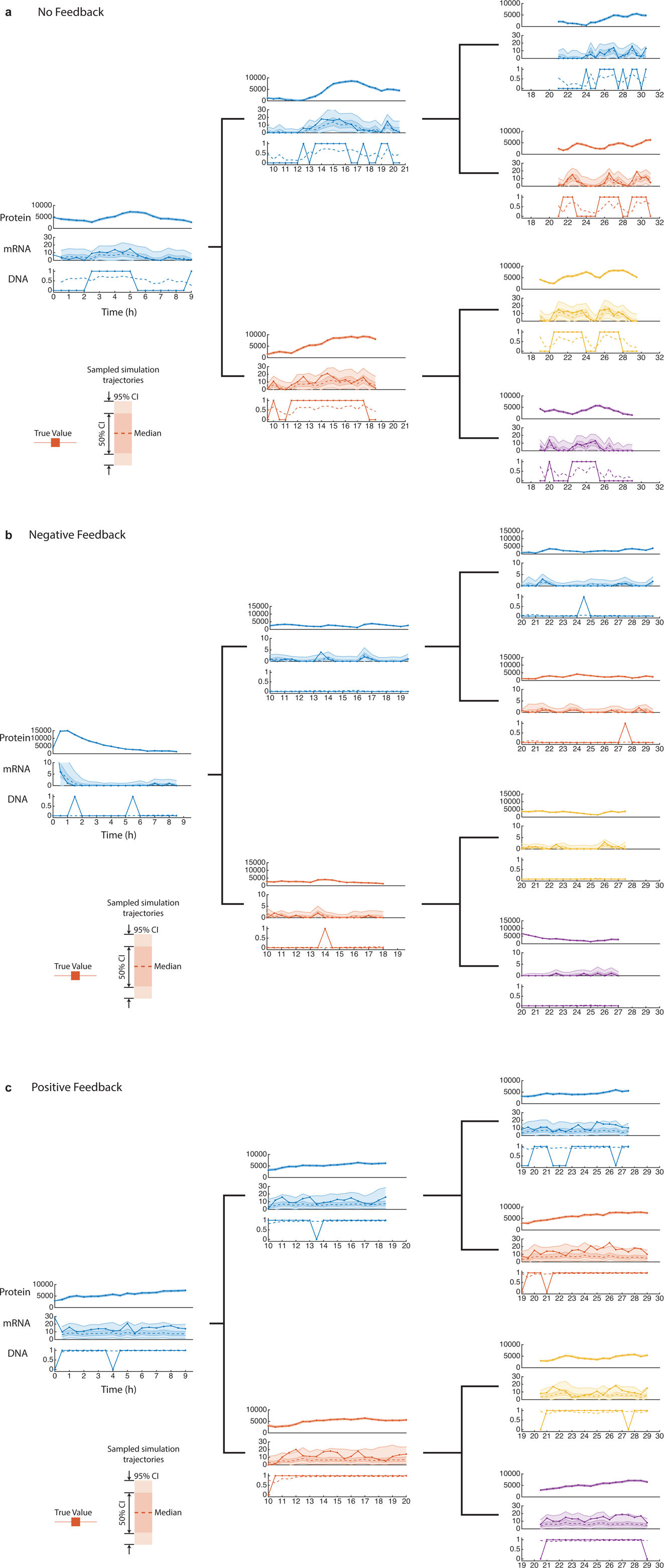
The particle filter provides estimates latent mRNA and DNA trajectories. Protein (top), mRNA (middle) and DNA (bottom) trajectories are estimated by STILT and show very good agreement between the true value (solid) and the resampled trajectories. The 50% and 95% confidence intervals of the resampled trajectories are shown as band plots, along with the median (dashed line). For DNA the true value (solid) is compared with the mean over the sampled histories (dashed). The trajectories for each cell of the simulated lineage are plotted separately for improved visibility. In each case the correct model is assumed for the simulated data set. Results are shown for the (A) No Feedback, (B) Negative Feedback and (C) Positive Feedback models.

**Figure S3:**
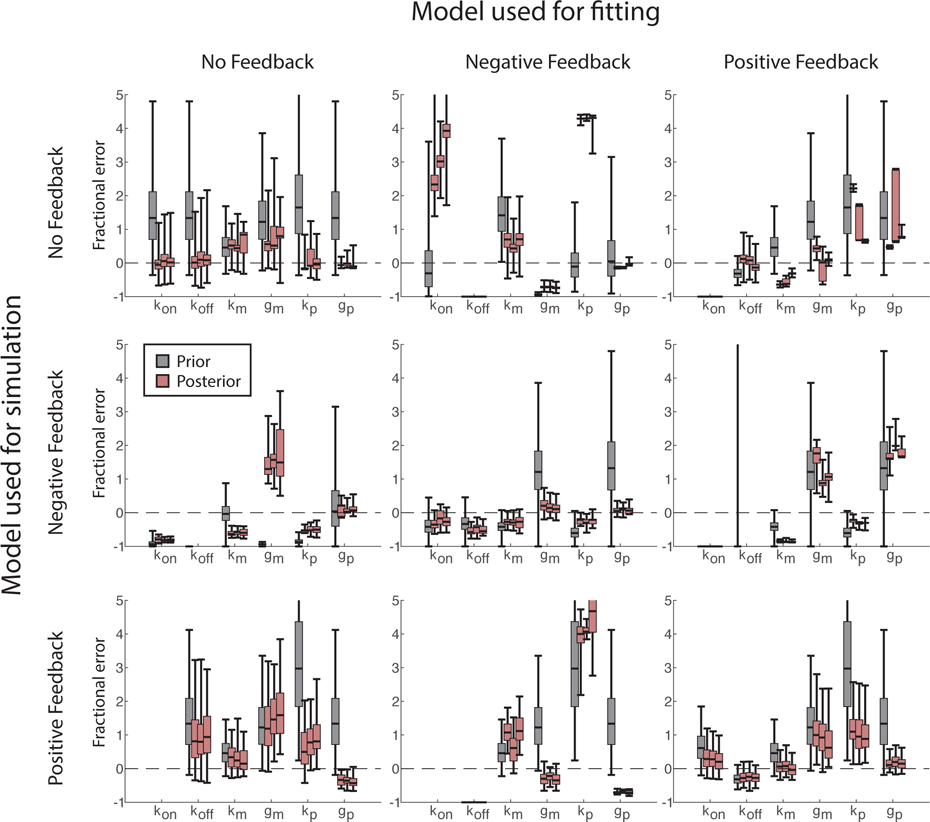
Posterior distributions of model parameters are robustly estimated in most cases. We estimate the posterior distribution of each model parameter for every combination of model used for data generation and model used for inference. Each combination is fit three times (red, 95% confidence interval) to provide an estimation of the robustness of the inference procedure, and compared against the prior distribution (gray). (A) Data generated with the No Feedback model. (B) Data generated with the Negative Feedback. (C) Data generated using the Positive Feedback model.

**Figure S4:**
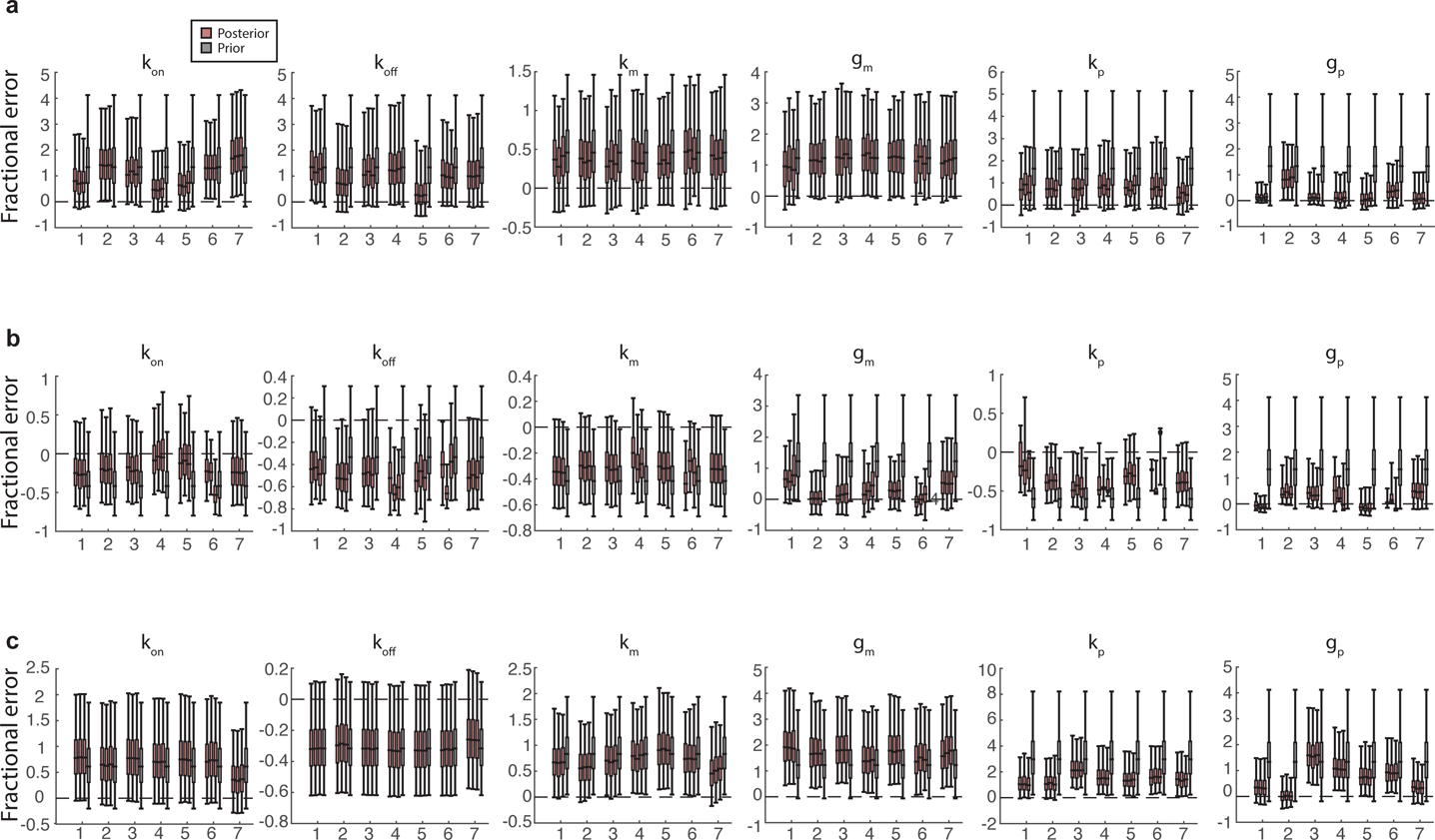
Parameter inference without including cellular lineage trees fails to converge to the correct model parameters. Single-cell-based inference was performed for each of the seven cells from each of the three synthetic datasets. The fractional error of each posterior is plotted and compared to the prior for each model parameter and dataset. The posteriors are robustly estimated, across the repeats of the inference procedure. However, STILT’s tree-based inference shows improved parameter convergence to the true value compared to the single-cell-based inference for most parameters. Additionally, there is substantial variation among cells due to the difference in information content of their respective trajectories. (A) No Feedback, (B) Negative Feedback, and (C) Positive Feedback.

**Figure S5:**
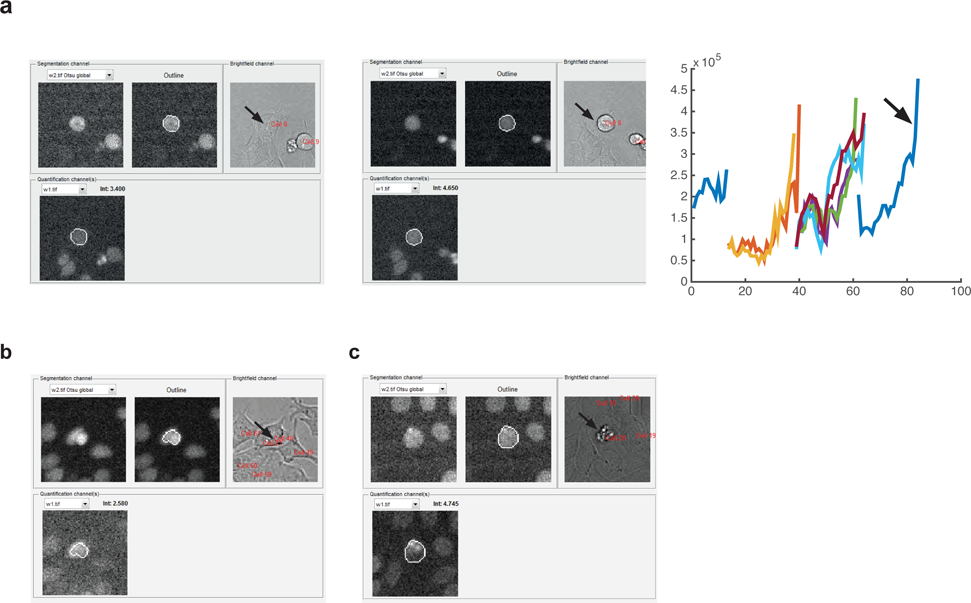
We pre-process quantified fluorescence lineage trees to remove inaccurate data. (A) Many cells show a substantial change in morphology, going from a flat (left) to round (middle) morphology just prior to division. This change often leads to an artificial jump in the quantified fluorescence intensity (right) at the last time point prior to division in many cells, and is thus censored. (B) Fluorescence signal at time points where cell nuclei overlap cannot be reliably quantified and are also censored. (C) Time points showing large fluctuations in fluorescence intensity due to contamination overlapping the segmented nuclei are likewise excluded.

**Figure S6:**
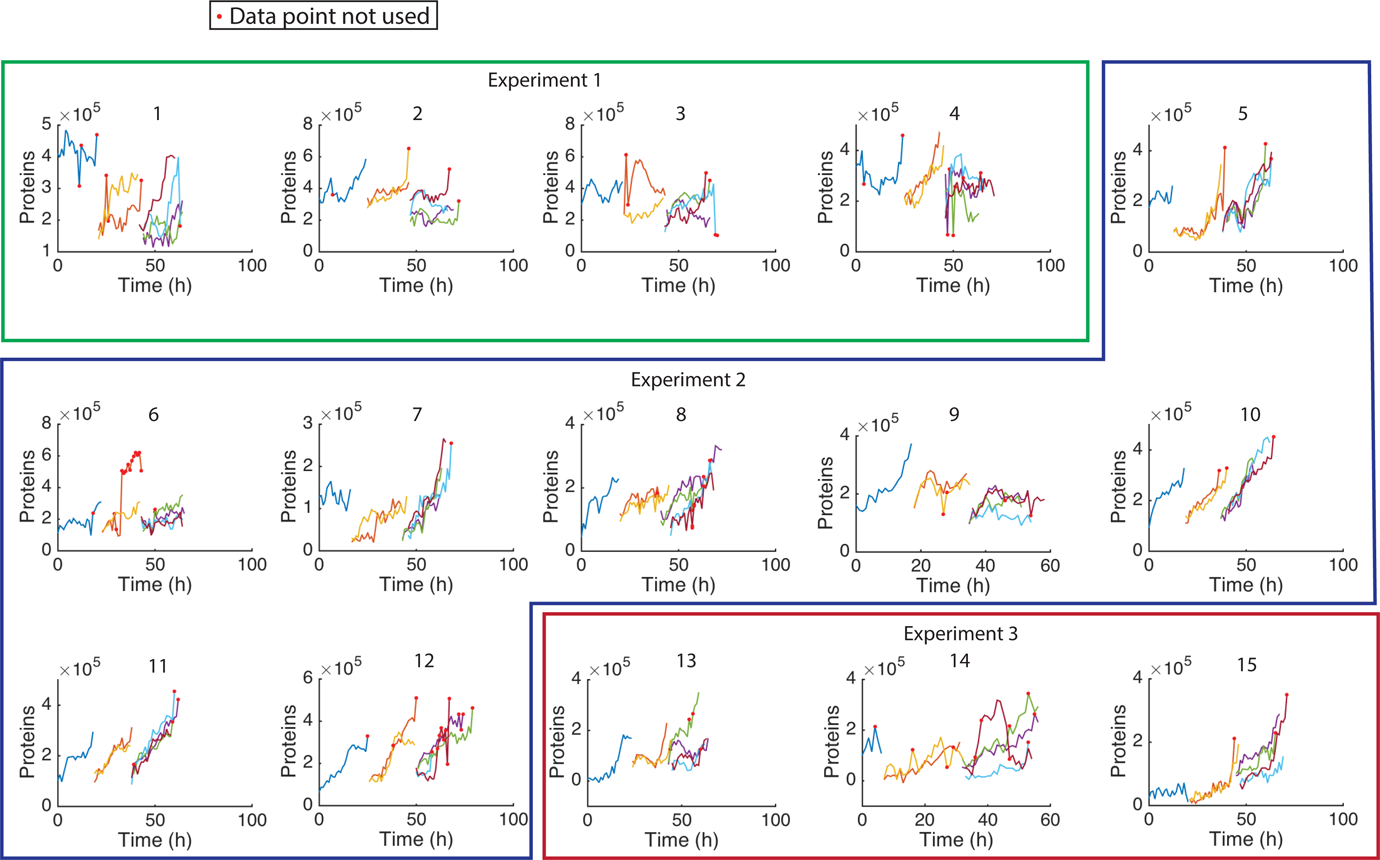
We obtain 15 NanogVENUS subtrees of 7 cells from 3 experiments. 15 NanogVENUS subtrees of 7 cells each are fitted with each of the autoregulatory models. The subtrees are obtained from 7 microscopy positions from 3 independent experiments, obtained by dividing trees into non-overlapping subtrees of seven immediately-related cells. Unreliable data points (red) are censored (see Methods) and not used for inference.

**Figure S7:**
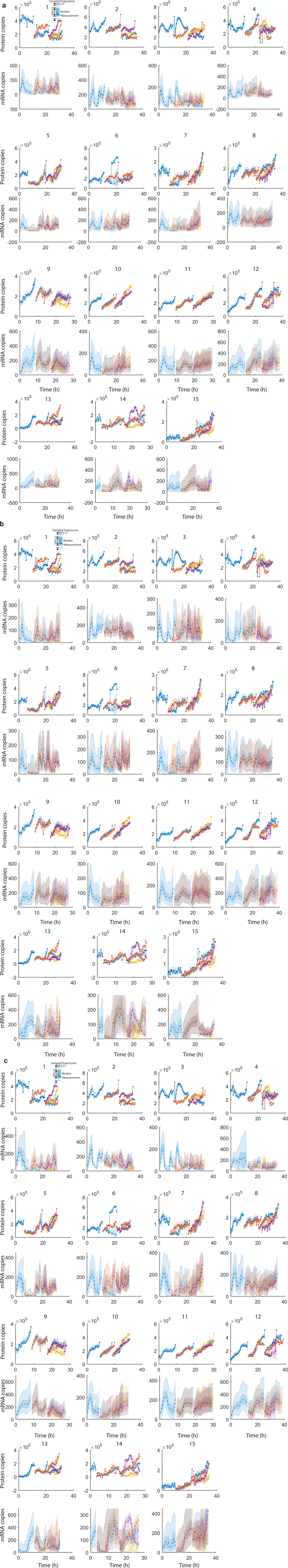
The particle filter samples from latent trajectories for the protein (top) and mRNA (bottom) conditional on the observed protein measurements. The particle filter was performed for each of the NanogVENUS subtrees using the (A) No Feedback, (B) Negative Feedback and (C) Positive Feedback models.

**Figure S8:**
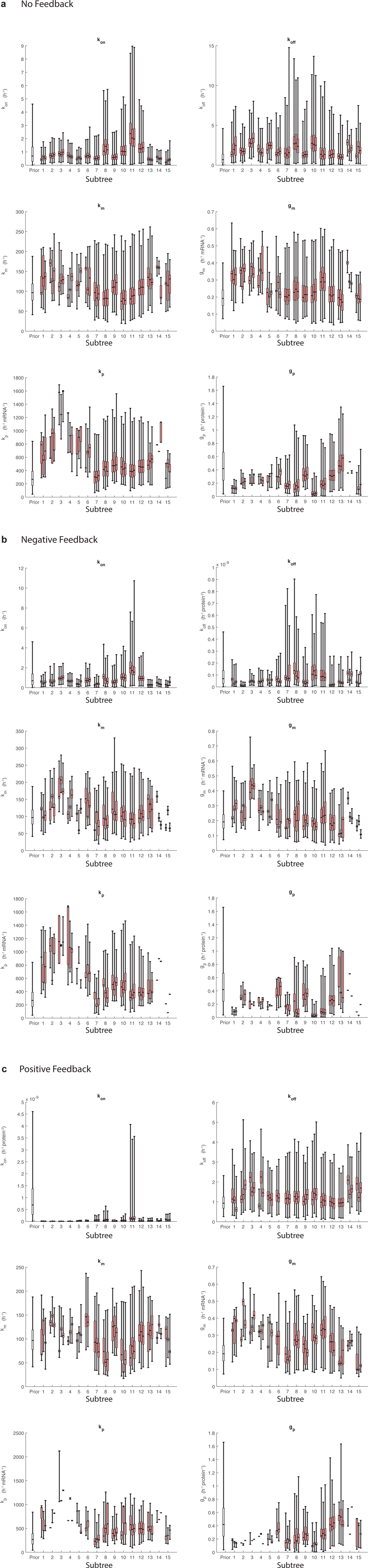
We estimate the posterior distribution of all model parameters for every NanogVENUS subtree, fit with each of the three autoregulatory models. Each combination is performed in triplicate (red) to assess robustness with respect to the stochastic nature of the inference procedure. Priors distributions are shown in gray. Each subtree was fit with the (A) No Feedback, (B) Negative Feedback and (C) Positive Feedback models.

**Figure S9:**
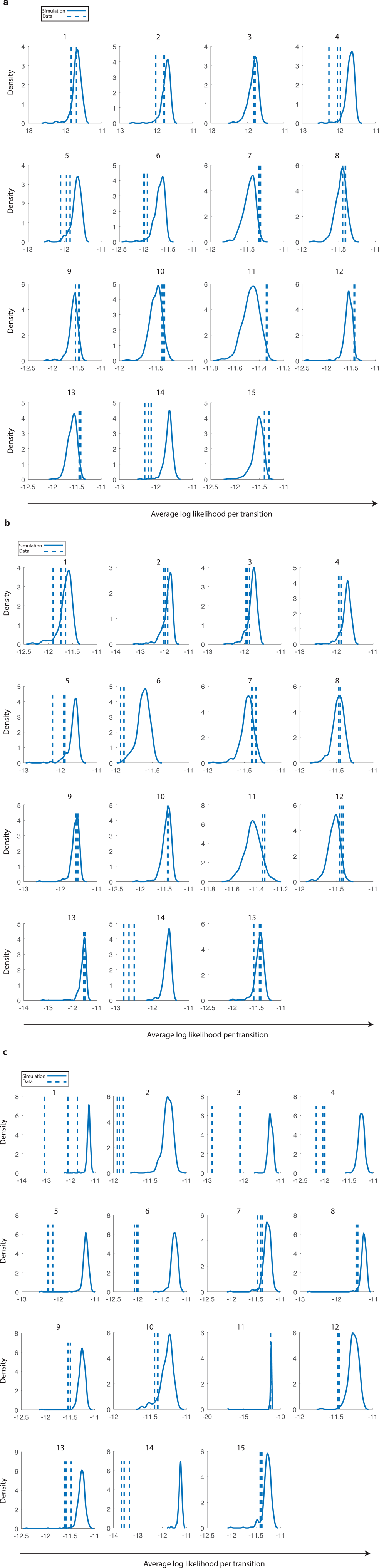
We use the goodness-of-fit test to determine which auto-regulatory models are compatible with which Nanog subtrees. We compute the goodness-of-fit quantiles of the real data with respect to simulations for the (A) No Feedback model, (B) Negative Feedback model and (C)Positive Feedback model.

**Figure S10:**
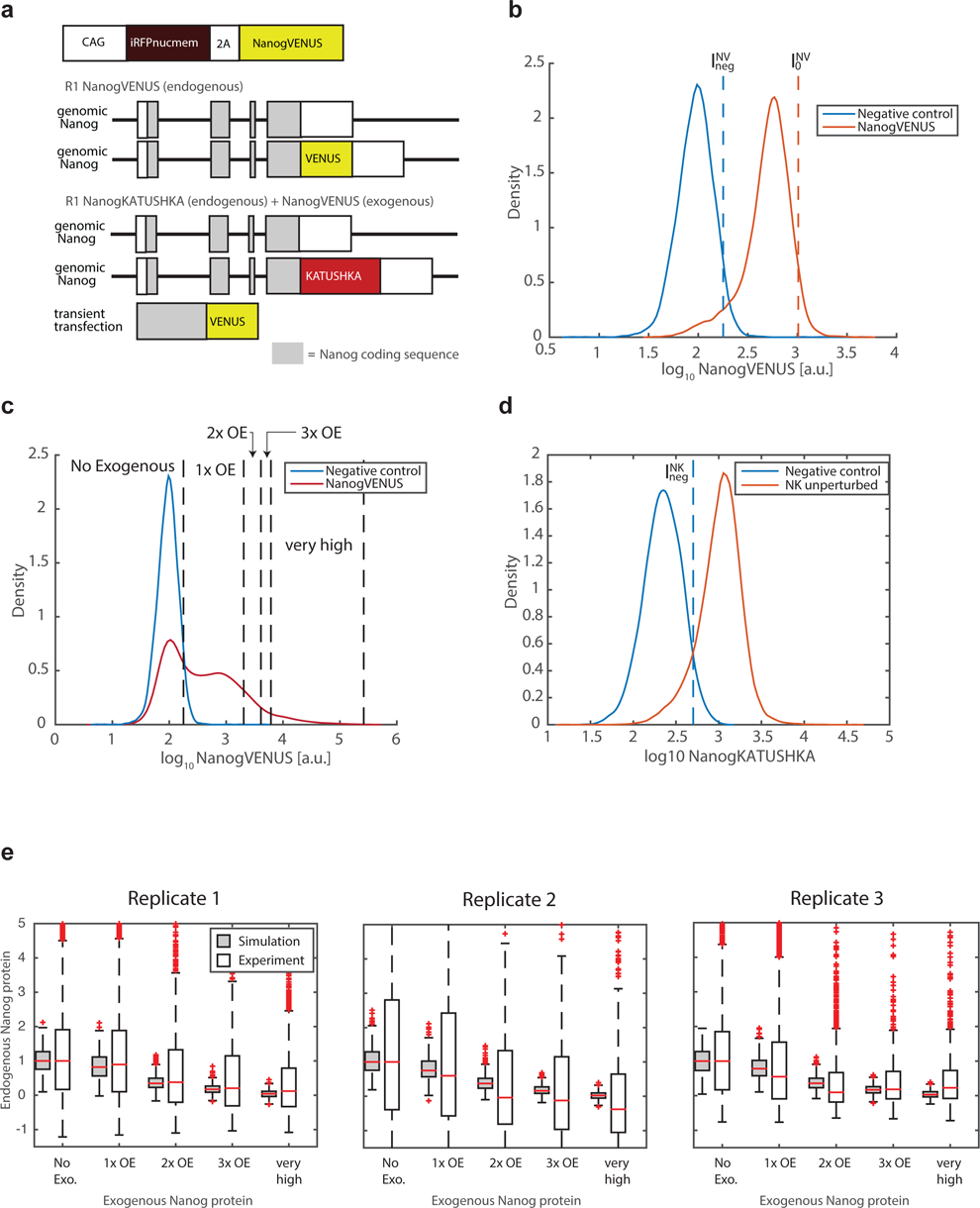
We estimate autofluorescence and expression levels of endogenous Nanog reporters. (A) Top: Schematic Nanog overexpression construct. The NanogVENUS and iRFPnucmem proteins are translated at similar levels. iRFPnucmem allows direct comparison of the construct with an empty vector (no NanogVENUS) and Nanog alone (without VENUS). CAG = constitutive promoter; 2A = sequence for protein “cleavage”; iRFP (targeted to nuclear membrane) and VENUS = fluorescent proteins. Bottom: Schematic representation of cell lines. Cells express a fluorescently labeled Nanog protein (either VENUS or KATUSHKA) from one endogenous allele. Exogenous NanogVENUS was transiently transfected 2 days before analysis. (B) We estimate the autofluorescence of NanogVENUS (I_neg_^NV^) using wild-type mESCs. We also estimate the 0.95 quantile of endogenous NanogVENUS (I_0_^NV^) in unperturbed NV cells. (C) We measure NanogVENUS levels and define compartments of exogenous Nanog as No Exogenous (below negative control levels); 1x, 2x, and 3x overexpression corresponding to up to 100%, 200% and 300%, respectively, of wild-type Nanog levels; and very high which has more than 300% wild-type Nanog expression. (D) We estimate the autofluorescence level (0.95 quantile) of NanogKATUSHKA (I_neg_^NK^) using the NanogVENUS cell line. The intensity distribution of unperturbed NanogKATUSHKA mESCs is shown for comparison. (E) We compare simulated (gray) and measured (white) down-regulation relative to the median of the No Exogenous overexpression compartment, for three experimental replicates. Distributions are of the relative expression of individual cells for both experiment and simulation.

## Supplementary Table captions

**Table S1:** Model parameters used for generating the synthetic dataset. Parameters differ slightly between the No Feedback, Negative Feedback, Positive Feedback models so as to provide similar order of magnitude between trajectories generated from each model. The mRNA degradation (g_m_) rate constants have units of per hour per count mRNA, and the protein degradation rate (g_p_) and translation (k_p_) per hour per count protein.

| Parameter | Units | No Feedback | Negative Feedback | Positive Feedback |
| --- | --- | --- | --- | --- |
| $k_{\text{on}}$ | $\text{h}^{-1}$ | 1 | 10 | $5 \times 10^{-7} \text{ protein}^{-2}$ |
| $k_{\text{off}}$ | $\text{h}^{-1}$ | 1 | $10^{-4} \text{ protein}^{-2}$ | 1 |
| $k_{\text{m}}$ | $\text{h}^{-1}$ | 40 | 100 | 40 |
| $g_{\text{m}}$ | $\text{h}^{-1} \text{ mRNA}^{-1}$ | 3 | 3 | 3 |
| $k_{\text{p}}$ | $\text{h}^{-1} \text{ mRNA}^{-1}$ | 300 | 2000 | 200 |
| $g_{\text{p}}$ | $\text{h}^{-1} \text{ protein}^{-1}$ | 0.4 | 0.4 | 0.4 |

**Table S2:**
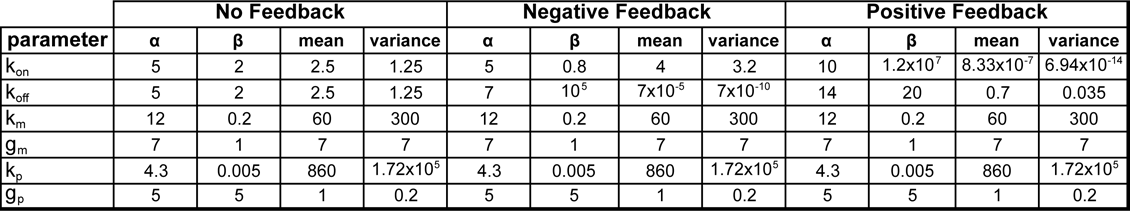
Parameters of the gamma prior distributions Γ (α, β) used for inference of the synthetic datasets.

**Table S3:**
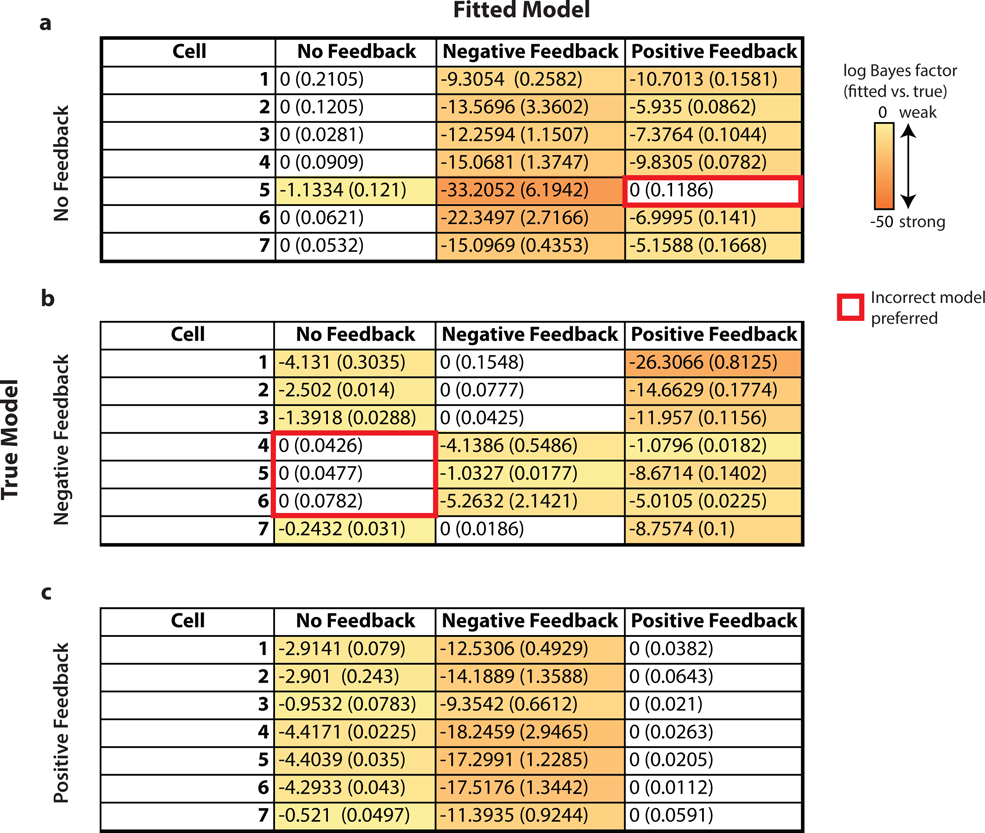
Bayes Factor analysis shows that model selection is less robust for single-cell-based inference. The computed log Bayes Factors of each model indicate that the incorrect model is preferred (red boxes) for one cell in the (A) No Feedback dataset, and for 3 cells in the (B) Negative Feedback dataset. (C) The Positive Feedback model is inferred correctly for each of the Positive Feedback datasets. Each row indicates the difference of the marginal log likelihood (mean, s.d.; n=3) of each fitted model and the marginal log likelihood of the best model for that cell/dataset, i.e. a value of 0 indicates the best model for that cell/dataset.

**Table S4:** Parameters characterizing the gamma distributions used for parameter Γ (α, β) prior distributions for the NanogVENUS subtrees inference.

| parameter | No Feedback |  |  |  | Negative Feedback |  |  |  | Positive Feedback |  |  |  |
| --- | --- | --- | --- | --- | --- | --- | --- | --- | --- | --- | --- | --- |
| | $\alpha$ | $\beta$ | mean | variance | $\alpha$ | $\beta$ | mean | variance | $\alpha$ | $\beta$ | mean | variance |
| $k_{on}$ | 1 | 1 | 1 | 1 | 1 | 1 | 1 | 1 | 1 | 1.00E+09 | 1.00E-09 | 1.00E-18 |
| $k_{off}$ | 1 | 1 | 1 | 1 | 1 | 1.00E+10 | 1.00E-10 | 1.00E-20 | 5 | 5 | 1 | 0.2 |
| $k_m$ | 10 | 0.1 | 100 | 1000 | 10 | 0.1 | 100 | 1000 | 10 | 0.1 | 100 | 1000 |
| $g_m$ | 8 | 40 | 0.2 | 0.005 | 8 | 40 | 0.2 | 0.005 | 8 | 40 | 0.2 | 0.005 |
| $k_p$ | 3 | 0.01 | 300 | 30000 | 3 | 0.01 | 300 | 30000 | 3 | 0.01 | 300 | 30000 |
| $g_p$ | 2 | 4 | 0.5 | 0.125 | 2 | 4 | 0.5 | 0.125 | 2 | 4 | 0.5 | 0.125 |

**Table S5:**
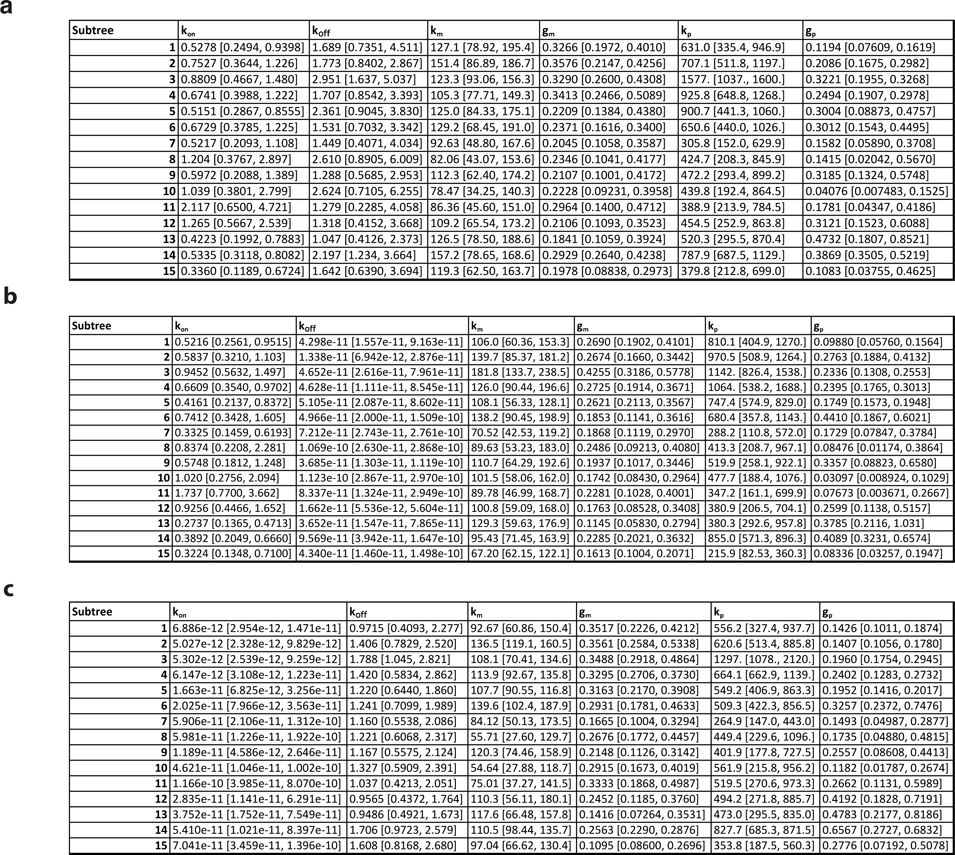
We estimate the median (95% confidence interval) of the posterior distributions of each model parameter for every NanogVENUS subtree using the (A) No Feedback, (B) Negative Feedback, and (C) Positive Feedback models.

| Subtree | k<sub>on</sub> | k<sub>off</sub> | k<sub>m</sub> | g<sub>m</sub> | k<sub>p</sub> | g<sub>p</sub> |
| --- | --- | --- | --- | --- | --- | --- |
| 1 | 0.5278 [0.2494, 0.9398] | 1.689 [0.7351, 4.511] | 127.1 [78.92, 195.4] | 0.3266 [0.1972, 0.4010] | 631.0 [335.4, 946.9] | 0.1194 [0.07609, 0.1619] |
| 2 | 0.7527 [0.3644, 1.226] | 1.773 [0.8402, 2.867] | 151.4 [86.89, 186.7] | 0.3576 [0.2147, 0.4256] | 707.1 [511.8, 1197.] | 0.2086 [0.1675, 0.2982] |
| 3 | 0.8809 [0.4667, 1.480] | 2.951 [1.637, 5.037] | 123.3 [93.06, 156.3] | 0.3290 [0.2600, 0.4308] | 1577. [1037., 1600.] | 0.3221 [0.1955, 0.3268] |
| 4 | 0.6741 [0.3988, 1.222] | 1.707 [0.8542, 3.393] | 105.3 [77.71, 149.3] | 0.3413 [0.2466, 0.5089] | 925.8 [648.8, 1268.] | 0.2494 [0.1907, 0.2978] |
| 5 | 0.5151 [0.2867, 0.8555] | 2.361 [0.9045, 3.830] | 125.0 [84.33, 175.1] | 0.2209 [0.1384, 0.4380] | 900.7 [441.3, 1060.] | 0.3004 [0.08873, 0.4757] |
| 6 | 0.6729 [0.3785, 1.225] | 1.531 [0.7032, 3.342] | 129.2 [68.45, 191.0] | 0.2371 [0.1616, 0.3400] | 650.6 [440.0, 1026.] | 0.3012 [0.1543, 0.4495] |
| 7 | 0.5217 [0.2093, 1.108] | 1.449 [0.4071, 4.034] | 92.63 [48.80, 167.6] | 0.2045 [0.1058, 0.3587] | 305.8 [152.0, 629.9] | 0.1582 [0.05890, 0.3708] |
| 8 | 1.204 [0.3767, 2.897] | 2.610 [0.8905, 6.009] | 82.06 [43.07, 153.6] | 0.2346 [0.1041, 0.4177] | 424.7 [208.3, 845.9] | 0.1415 [0.02042, 0.5670] |
| 9 | 0.5972 [0.2088, 1.389] | 1.288 [0.5685, 2.953] | 112.3 [62.40, 174.2] | 0.2107 [0.1001, 0.4172] | 472.2 [293.4, 899.2] | 0.3185 [0.1324, 0.5748] |
| 10 | 1.039 [0.3801, 2.799] | 2.624 [0.7105, 6.255] | 78.47 [34.25, 140.3] | 0.2228 [0.09231, 0.3958] | 439.8 [192.4, 864.5] | 0.04076 [0.007483, 0.1525] |
| 11 | 2.117 [0.6500, 4.721] | 1.279 [0.2285, 4.058] | 86.36 [45.60, 151.0] | 0.2964 [0.1400, 0.4712] | 388.9 [213.9, 784.5] | 0.1781 [0.04347, 0.4186] |
| 12 | 1.265 [0.5667, 2.539] | 1.318 [0.4152, 3.668] | 109.2 [65.54, 173.2] | 0.2106 [0.1093, 0.3523] | 454.5 [252.9, 863.8] | 0.3121 [0.1523, 0.6088] |
| 13 | 0.4223 [0.1992, 0.7883] | 1.047 [0.4126, 2.373] | 126.5 [78.50, 188.6] | 0.1841 [0.1059, 0.3924] | 520.3 [295.5, 870.4] | 0.4732 [0.1807, 0.8521] |
| 14 | 0.5335 [0.3118, 0.8082] | 2.197 [1.234, 3.664] | 157.2 [78.65, 168.6] | 0.2929 [0.2640, 0.4238] | 787.9 [687.5, 1129.] | 0.3869 [0.3505, 0.5219] |
| 15 | 0.3360 [0.1189, 0.6724] | 1.642 [0.6390, 3.694] | 119.3 [62.50, 163.7] | 0.1978 [0.08838, 0.2973] | 379.8 [212.8, 699.0] | 0.1083 [0.03755, 0.4625] |

b
| Subtree | k<sub>on</sub> | k<sub>off</sub> | k<sub>m</sub> | g<sub>m</sub> | k<sub>p</sub> | g<sub>p</sub> |
| --- | --- | --- | --- | --- | --- | --- |
| 1 | 0.5216 [0.2561, 0.9515] | 4.298e-11 [1.557e-11, 9.163e-11] | 106.0 [60.36, 153.3] | 0.2690 [0.1902, 0.4101] | 810.1 [404.9, 1270.] | 0.09880 [0.05760, 0.1564] |
| 2 | 0.5837 [0.3210, 1.103] | 1.338e-11 [6.942e-12, 2.876e-11] | 139.7 [85.37, 181.2] | 0.2674 [0.1660, 0.3442] | 970.5 [508.9, 1264.] | 0.2763 [0.1884, 0.4132] |
| 3 | 0.9452 [0.5632, 1.497] | 4.652e-11 [2.616e-11, 7.961e-11] | 181.8 [133.7, 238.5] | 0.4255 [0.3186, 0.5778] | 1142. [826.4, 1538.] | 0.2336 [0.1308, 0.2553] |
| 4 | 0.6609 [0.3540, 0.9702] | 4.628e-11 [1.111e-11, 8.545e-11] | 126.0 [90.44, 196.6] | 0.2725 [0.1914, 0.3671] | 1064. [538.2, 1688.] | 0.2395 [0.1765, 0.3013] |
| 5 | 0.4161 [0.2137, 0.8372] | 5.105e-11 [2.087e-11, 8.602e-11] | 108.1 [56.33, 128.1] | 0.2621 [0.2113, 0.3567] | 747.4 [574.9, 829.0] | 0.1749 [0.1573, 0.1948] |
| 6 | 0.7412 [0.3428, 1.605] | 4.966e-11 [2.000e-11, 1.509e-10] | 138.2 [90.45, 198.9] | 0.1853 [0.1141, 0.3616] | 680.4 [357.8, 1143.] | 0.4410 [0.1867, 0.6021] |
| 7 | 0.3325 [0.1459, 0.6193] | 7.212e-11 [2.743e-11, 2.761e-10] | 70.52 [42.53, 119.2] | 0.1868 [0.1119, 0.2970] | 288.2 [110.8, 572.0] | 0.1729 [0.07847, 0.3784] |
| 8 | 0.8374 [0.2208, 2.281] | 1.069e-10 [2.630e-11, 2.868e-10] | 89.63 [53.23, 183.0] | 0.2486 [0.09213, 0.4080] | 413.3 [208.7, 967.1] | 0.08476 [0.01174, 0.3864] |
| 9 | 0.5748 [0.1812, 1.248] | 3.685e-11 [1.303e-11, 1.119e-10] | 110.7 [64.29, 192.6] | 0.1937 [0.1017, 0.3446] | 519.9 [258.1, 922.1] | 0.3357 [0.08823, 0.6580] |
| 10 | 1.020 [0.2756, 2.094] | 1.123e-10 [2.867e-11, 2.970e-10] | 101.5 [58.06, 162.0] | 0.1742 [0.08430, 0.2964] | 477.7 [188.4, 1076.] | 0.03097 [0.008924, 0.1029] |
| 11 | 1.737 [0.7700, 3.662] | 8.337e-11 [1.324e-11, 2.949e-10] | 89.78 [46.99, 168.7] | 0.2281 [0.1028, 0.4001] | 347.2 [161.1, 699.9] | 0.07673 [0.003671, 0.2667] |
| 12 | 0.9256 [0.4466, 1.652] | 1.662e-11 [5.536e-12, 5.604e-11] | 100.8 [59.09, 168.0] | 0.1763 [0.08528, 0.3408] | 380.9 [206.5, 704.1] | 0.2599 [0.1138, 0.5157] |
| 13 | 0.2737 [0.1365, 0.4713] | 3.652e-11 [1.547e-11, 7.865e-11] | 129.3 [59.63, 176.9] | 0.1145 [0.05830, 0.2794] | 380.3 [292.6, 957.8] | 0.3785 [0.2116, 1.031] |
| 14 | 0.3892 [0.2049, 0.6660] | 9.569e-11 [3.942e-11, 1.647e-10] | 95.43 [71.45, 163.9] | 0.2285 [0.2021, 0.3632] | 855.0 [571.3, 896.3] | 0.4089 [0.3231, 0.6574] |
| 15 | 0.3224 [0.1348, 0.7100] | 4.340e-11 [1.460e-11, 1.498e-10] | 67.20 [62.15, 122.1] | 0.1613 [0.1004, 0.2071] | 215.9 [82.53, 360.3] | 0.08336 [0.03257, 0.1947] |

| Subtree | k<sub>on</sub> | k<sub>off</sub> | k<sub>m</sub> | g<sub>m</sub> | k<sub>p</sub> | g<sub>p</sub> |
| --- | --- | --- | --- | --- | --- | --- |
| 1 | 6.886e-12 [2.954e-12, 1.471e-11] | 0.9715 [0.4093, 2.277] | 92.67 [60.86, 150.4] | 0.3517 [0.2226, 0.4212] | 556.2 [327.4, 937.7] | 0.1426 [0.1011, 0.1874] |
| 2 | 5.027e-12 [2.328e-12, 9.829e-12] | 1.406 [0.7829, 2.520] | 136.5 [119.1, 160.5] | 0.3561 [0.2584, 0.5338] | 620.6 [513.4, 885.8] | 0.1407 [0.1056, 0.1780] |
| 3 | 5.302e-12 [2.539e-12, 9.259e-12] | 1.788 [1.045, 2.821] | 108.1 [70.41, 134.6] | 0.3488 [0.2918, 0.4864] | 1297. [1078., 2120.] | 0.1960 [0.1754, 0.2945] |
| 4 | 6.147e-12 [3.108e-12, 1.223e-11] | 1.420 [0.5834, 2.862] | 113.9 [92.67, 135.8] | 0.3295 [0.2706, 0.3730] | 664.1 [662.9, 1139.] | 0.2402 [0.1283, 0.2732] |
| 5 | 1.663e-11 [6.825e-12, 3.256e-11] | 1.220 [0.6440, 1.860] | 107.7 [90.55, 116.8] | 0.3163 [0.2170, 0.3908] | 549.2 [406.9, 863.3] | 0.1952 [0.1416, 0.2017] |
| 6 | 2.025e-11 [7.966e-12, 3.563e-11] | 1.241 [0.7099, 1.989] | 139.6 [102.4, 187.9] | 0.2931 [0.1781, 0.4633] | 509.3 [422.3, 856.5] | 0.3257 [0.2372, 0.7476] |
| 7 | 5.906e-11 [2.106e-11, 1.312e-10] | 1.160 [0.5538, 2.086] | 84.12 [50.13, 173.5] | 0.1665 [0.1004, 0.3294] | 264.9 [147.0, 443.0] | 0.1493 [0.04987, 0.2877] |
| 8 | 5.981e-11 [1.226e-11, 1.922e-10] | 1.221 [0.6068, 2.317] | 55.71 [27.60, 129.7] | 0.2676 [0.1772, 0.4457] | 449.4 [229.6, 1096.] | 0.1735 [0.04880, 0.4815] |
| 9 | 1.189e-11 [4.586e-12, 2.646e-11] | 1.167 [0.5575, 2.124] | 120.3 [74.46, 158.9] | 0.2148 [0.1126, 0.3142] | 401.9 [177.8, 727.5] | 0.2557 [0.08608, 0.4413] |
| 10 | 4.621e-11 [1.046e-11, 1.002e-10] | 1.327 [0.5909, 2.391] | 54.64 [27.88, 118.7] | 0.2915 [0.1673, 0.4019] | 561.9 [215.8, 956.2] | 0.1182 [0.01787, 0.2674] |
| 11 | 1.166e-10 [3.985e-11, 8.070e-10] | 1.037 [0.4213, 2.051] | 75.01 [37.27, 141.5] | 0.3333 [0.1868, 0.4987] | 519.5 [270.6, 973.3] | 0.2662 [0.1131, 0.5989] |
| 12 | 2.835e-11 [1.141e-11, 6.291e-11] | 0.9565 [0.4372, 1.764] | 110.3 [56.11, 180.1] | 0.2452 [0.1185, 0.3760] | 494.2 [271.8, 885.7] | 0.4192 [0.1828, 0.7191] |
| 13 | 3.752e-11 [1.752e-11, 7.549e-11] | 0.9486 [0.4921, 1.673] | 117.6 [66.48, 157.8] | 0.1416 [0.07264, 0.3531] | 473.0 [295.5, 835.0] | 0.4783 [0.2177, 0.8186] |
| 14 | 5.410e-11 [1.021e-11, 8.397e-11] | 1.706 [0.9723, 2.579] | 110.5 [98.44, 135.7] | 0.2563 [0.2290, 0.2876] | 827.7 [685.3, 871.5] | 0.6567 [0.2727, 0.6832] |
| 15 | 7.041e-11 [3.459e-11, 1.396e-10] | 1.608 [0.8168, 2.680] | 97.04 [66.62, 130.4] | 0.1095 [0.08600, 0.2696] | 353.8 [187.5, 560.3] | 0.2776 [0.07192, 0.5078] |

**Table S6:** Evidence of each subtree/model combination. We estimate the evidence (marginal log likelihood) of each model applied to each dataset using the particle filter (mean, s.e., n=3).

| **Subtree** | **No Feedback** | **Negative Feedback** | **Positive Feedback** |
| --- | --- | --- | --- |
| **1** | -1634.1 (0.4333) | -1636.3000 (0.9074) | -1641.6333 (0.8172) |
| **2** | -1814.7 (0.2646) | -1824.6000 (0.7211) | -1843.4667 (4.9320) |
| **3** | -1835.4 (0.5568) | -1834.6333 (1.1837) | -1865.3333 (4.3910) |
| **4** | -1700.3 (0.1528) | -1703.6333 (0.2906) | -1711.4000 (2.0075) |
| **5** | -1838.6 (0.4842) | -1843.9333 (1.5603) | -1848.7667 (2.1184) |
| **6** | -1597.1 (0.7126) | -1595.7000 (0.2000) | -1602.7858 (0.0231) |
| **7** | -1747.4 (0.1453) | -1753.1000 (0.7095) | -1754.5000 (0.1528) |
| **8** | -1820.4 (0.0882) | -1821.8667 (0.2603) | -1825.2333 (0.2333) |
| **9** | -1450.7 (0.0577) | -1454.8000 (0.1528) | -1453.7333 (0.2333) |
| **10** | -1554.4 (0.1732) | -1553.9000 (0.0000) | -1560.1000 (0.2517) |
| **11** | -1615.6 (0.0882) | -1615.1000 (0.0577) | -1617.6000 (0.1732) |
| **12** | -1591.0 (0.0577) | -1592.7333 (0.0667) | -1598.5333 (0.0882) |
| **13** | -1546.5 (0.1528) | -1551.8000 (0.1732) | -1554.8667 (0.6386) |
| **14** | -1714.6 (5.2129) | -1769.1000 (7.7253) | -1795.0667 (9.8660) |
| **15** | -1803.4 (0.4933) | -1832.0667 (0.2333) | -1810.9333 (0.7796) |

**Table S7:** We use the goodness-of-fit test to determine which autoregulatory models are compatible with the NanogVENUS subtrees. For each subtree and model we estimate the null distribution of the average marginal log likelihood per transition (see Figure S9). We use this empirical null distribution to compute the quantile of the average marginal log likelihood per transition of each dataset (mean, s.e.m., n=3 replicates). Each model is accepted if it is within the 95% confidence interval of the empirical distribution, marginally accepted if it is not accepted but within the 98% confidence interval, and rejected otherwise.

| Experiment | Subtree number | No Feedback | Negative Feedback | Positive Feedback |
| --- | --- | --- | --- | --- |
| 1 | 1 | 0.2978 (0.1125) | 0.2033 (0.0996) | 0.0033 (0.0033) |
|  | 2 | 0.1811 (0.0804) | 0.2578 (0.0617) | 0 (0) |
|  | 3 | 0.5478 (0.0473) | 0.1444 (0.0307) | 0 (0) |
|  | 4 | 0.0167 (0.0058) | 0.0944 (0.0195) | 0 (0) |
| 2 | 5 | 0.0567 (0.0291) | 0.0467 (0.0158) | 0 (0) |
|  | 6 | 0.0222 (0.004) | 0.0044 (0.0022) | 0 (0) |
|  | 7 | 0.9867 (0.0058) | 0.8478 (0.0478) | 0.0533 (0.019) |
|  | 8 | 0.7422 (0.0713) | 0.5322 (0.0356) | 0.0267 (0.0051) |
|  | 9 | 0.8389 (0.0879) | 0.6122 (0.0695) | 0.0067 (0) |
|  | 10 | 0.9044 (0.0292) | 0.59 (0.0386) | 0.0867 (0.0102) |
|  | 11 | 0.9956 (0.0011) | 0.9267 (0.0133) | 0.16 (0.0084) |
|  | 12 | 0.9756 (0.0048) | 0.9267 (0.0315) | 0.0078 (0.0011) |
| 3 | 13 | 0.97 (0.0133) | 0.6011 (0.083) | 0.02 (0.0033) |
|  | 14 | 0.0089 (0.0022) | 0 (0) | 0 (0) |
|  | 15 | 0.9789 (0.0211) | 0.3689 (0.1529) | 0.08 (0.0058) |

Reject ( $p < 0.01$ or $p > 0.99$ )Marginal ( $p < 0.025$ or $p > 0.975$ )
Accept

## Online Methods to: Exact Bayesian lineage tree-based inference identifies Nanog negative autoregulation in mouse embryonic stem cells

### 1 Chemical reaction networks

We consider the case of parameter inference and model comparison for stochastic models of gene regulation described by chemical reaction networks. A chemical reaction network consists of a set of chemical species (e.g. DNA, mRNA, protein, etc.) which may interact via a set of chemical reactions corresponding to synthesis, destruction, or modification.

Each reaction is defined by its stoichiometry, i.e. the quantity of each educt consumed and product produced by the reaction, and reaction rate. Reactions are presumed to take place stochastically as a function of the state of the system, i.e. the number of molecules of each species at a given point in time. The probability of a reaction occurring in infinitesimal time, called the reaction propensity, depends on the number of molecules of each educt available, the reaction volume (i.e. of the cellular compartment wherein the reaction takes place), and on a reaction constant. We consider reactions that are at most bimolecular, since reactions involving more than two species rarely occur in biology. Zeroth order reactions involve the production of a species with no dependence on an educt, for example due to constitutive production; their reaction propensities are constant. First order reactions proceed with propensity proportional to the number of available molecules of a single educt. Second order reactions involve two species, and their propensity is proportional to the abundances of both involved educts, and so on. The reaction constants *k* depend on the chemical species involved, temperature and reaction system volume. Reaction propensities are summarized in Table A1.

**Table A1:**
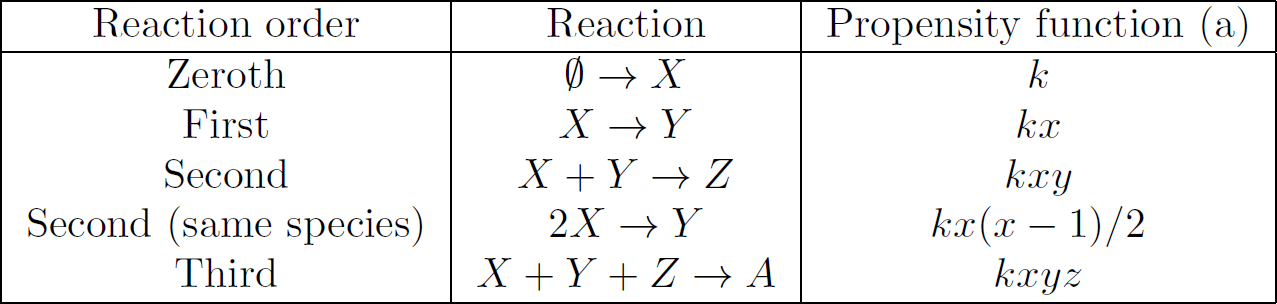
Reaction propensities for chemical reactions of zeroth, first, and second order. *k* denotes a kinetic constant and *x, y, z* the number of molecules of species *X, Y* and *Z*, respectively.

| Reaction order | Reaction | Propensity function (a) |
| --- | --- | --- |
| Zeroth | $\emptyset \rightarrow X$ | $k$ |
| First | $X \rightarrow Y$ | $kx$ |
| Second | $X + Y \rightarrow Z$ | $kxy$ |
| Second (same species) | $2X \rightarrow Y$ | $kx(x-1)/2$ |
| Third | $X + Y + Z \rightarrow A$ | $kxyz$ |

### 2 Inference of latent history and model parameters

#### 2.1 Inference using bootstrap particle filtering

STILT builds upon the recursive, simulation-based particle filter first introduced by Pitt and Sheppard [1]. The particle filter approximates the posterior distribution of the latent history of all chemical species for each cell, and all model parameters by iteratively including new observations.

Consider a chemical reaction network with *N_s_* chemical species, of which *N_o_* ≤ *N_s_* are observed via measurement. The vector of reaction constants governing the reactions of this network is denoted by *θ*. We denote by 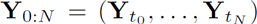 the set of observations obtained at a series of *N* discrete measurement time points *t*_0_,…, *t_N_*. Each observation 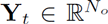 consists of measurements of *N_o_* observed chemical species. We assume that the observations **Y**_*t*_ constitute noisy measurements of the true unknown state of the system at time *t*, denoted by **X**_*t*_. We denote by 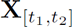 the path (trajectory) of the random variable **X**_*t*_ from time *t*_1_ to time *t*_2_, and denote by 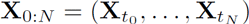 the value of **X**_*t*_ at the measurement time points *t*_0_,…, *t_N_*.

The objective of the bootstrap particle filter is to sample from the posterior joint density 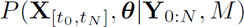 of latent trajectories 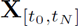 and parameters *θ* for a model with index *M*, given the observed data Y_0:*N*_. We drop the model index *M* for simplicity; when comparing models we will again introduce this notation. The latent trajectories 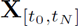 are realizations of a stochastic process, and the data **Y**_0:*N*_ represent noisy observations of (a function of) the latent process obtained at discrete times. The posterior joint density depends on the likelihood 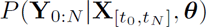 and parameter prior probability distribution *π*(*θ*) according to Bayes’ Law:

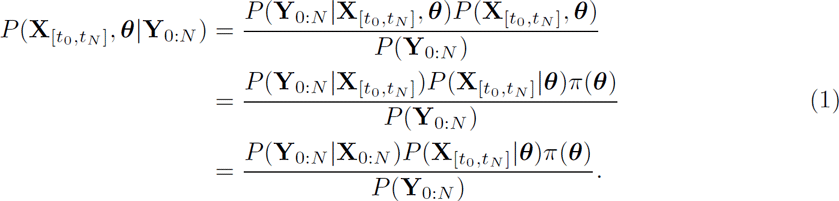

The simplification on the right side of (1) is possible since the probability of observing data **Y**_0:*N*_ given a latent trajectory 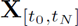 depends only on its value at the measurement timepoints **X**_0:*N*_. Furthermore, it does not depend on the underlying parameters *θ* of the stochastic process (measurement error is considered separately). The probability 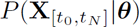 captures the evolution of the stochastic processes parameterized by *θ*.

The observations **Y**_0:*N*_ is related to 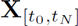 by a measurement function *g* with parameter *η*: 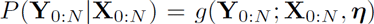. The function *g* depends on the measurement process and/or apparatus. For example, *g* might be a (multivariate) Gaussian in which case *η* contains the variance of the measurement process and potentially a scaling factor. We restrict ourselves to chemical reaction networks for which the stochastic process **X**_*t*_ is a Markov jump process on a subset of the integer lattice 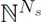, corresponding to molecular copy numbers reachable by the chemical reactions of the network. For such a system, the exact likelihood of the latent trajectory 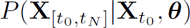, *θ* can be computed [2], and exact samples of 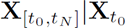 can be generated e.g. using Gillespie’s algorithm [3]. Note that the transition density (i.e. 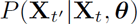, *t′* > *t*) of the stochastic process is in general not known, but can be approximated for small systems, e.g. using the Finite State Projection [4].

Assuming uncorrelated errors in the observation function *g*, the likelihood 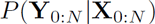 factorizes as:

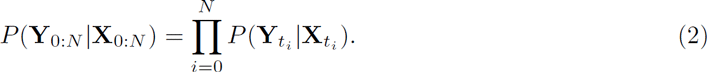

for a series of *N* observations.

Furthermore, the stochastic process **X**_*t*_ is Markovian such that the probability 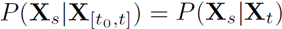 for some *s* ≥ *t*. The likelihood of the trajectory 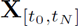 therefore decomposes as:

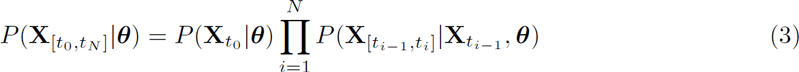

Following the derivation of Gordon *et al.* [5], we combine (2) and (3), and substitute into (1), to obtain a new expression for the posterior density:

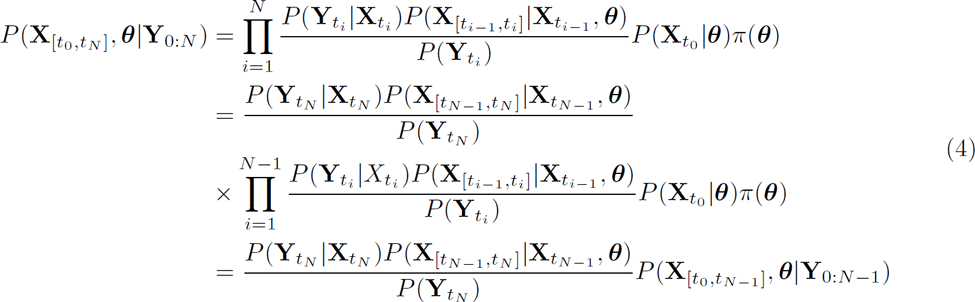

This can be rewritten as

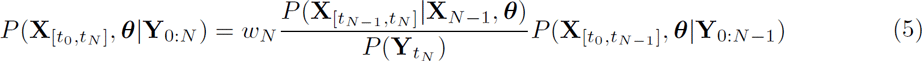

where 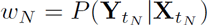.

Hence, there is a simple update rule relating the posterior distribution using observations until timepoint *t_N−_*_1_ to the posterior distribution with the next observation at timepoint *t_N_*. We note also that one can generate a sample from the posterior joint density at time *t_i_*, 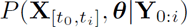, by first sampling a trajectory from the marginal distibution 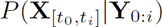 and then sampling a parameter 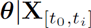, suggesting a Gibbs sampling approach.

These observations and the recursive factorization of the joint posterior (5) motivates the so-called bootstrap (recursive) particle filter [5], which iteratively generates samples (particles) from the posterior distribution conditioned on all prior observations:

#### Algorithm 1: Bootstrap particle filter

**Data:** A set of observed data points **Y**_0:*N*_ at timepoints *t*_0_,…, *t_N_*, parameter prior 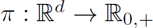, observation function *g*(**Y**; **X**, *η*) = *P* (**Y**, **X**), number of particles *K*

**Result:** A set of particles 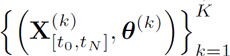 sampled from the posterior density 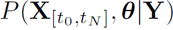

Initialization;
**for** *k=1… K* **do**

Sample parameter values from the prior: *θ*^(*k*)^ ~ *π*(*θ*);
Sample initial state conditional on first observed data point: 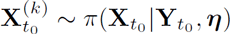;
Initialize particle weight to 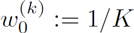
Iterate over observation time points;
**for** *i=1… N* **do**

Generate a set of particle indices *ε*^(*k*)^ ∈ {1,…, *K*}, *k* = 1,…, *K* such that
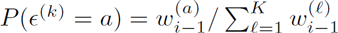
**for** *k* = 1… *K* **do**

Generate a sample trajectory 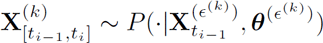;
Concatenate to previously sampled trajectory: 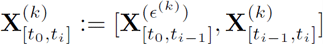;
Set the weight of the *k*^th^ particle to the likelihood: 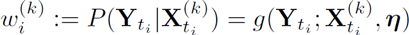;
Generate a new set of parameters *θ*^(*k*)^ from the conditional density: 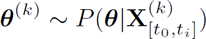;
Sample from the posterior;
Generate a set of particle indices *ε*^(*k*)^ ∈ {1,…, *K*}, *k* = 1,…, *K* such that
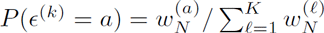;
Construct a sample of *K* particles from the posterior: 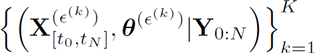

The recursive particle filter begins by sampling parameters *θ* from the parameter prior distribution *π*(*θ*), and an initial state 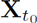 from its prior, for an ensemble of *K* particles, i.e. each particle is a sample from the joint density of 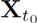 and *θ*. At each iteration *i*, the particles are resampled according to their normalized weights 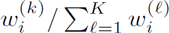, such that particles that have a state 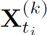 for which the current observation 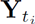 is likely are sampled more frequently. After updating the latent histories 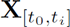, new samples are generated for the parameters conditional on those latent histories using a Gibbs sampling approach. Together the sample (**X**[*t_i_*]^(*k*)^, *θ*^(*k*)^) is used to simulate a new trajectory on the interval [*t_i_, t_i_*_+1_] using the stochastic simulation algorithm or variants [6, 7]. The result of the recursive particle filter at iteration *i* is an exact sample from the posterior joint density of 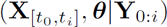, as shown in (4).

#### 2.2 Gamma priors

We consider the case of chemical reaction networks, in which case each parameter *θ*_1_,…, *θ_d_* ∈ *θ* corresponds to the kinetic constant of a chemical reaction (see Table A1). The inference procedure is significantly simplified if one assumes that the prior of each parameter is gamma distributed, and that the prior distributions of all parameters are conditionally independent:

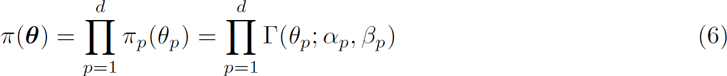

where *α_p_* and *β_p_* are the hyperparameters for the distribution of *θ_p_*, *p* = 1… *d*, and 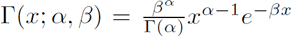. The two parameters *α* and *β* of the gamma distribution can be chosen e.g. to match a target mean *μ* and variance *σ*^2^:

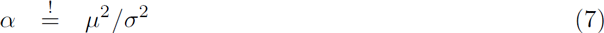

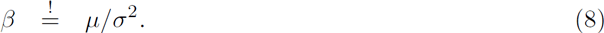

The assumption of conditional independence between model parameters is often justified, as information about the covariance of biological constants is often not available.

Using gamma priors for each model parameter, the likelihood *P* (**X**_[*t,t*+*τ*]_*|θ*) of a particular (fully-observed) realization of the Markov jump process **X**_*t*_ on the interval [*t, t* + *τ*] is conjugate to the prior, such that the conditional density *P* (*θ |***X**_[*t,t*+*τ*]_) is also gamma distributed (see Wilkinson *et al.* [2], p. 281):

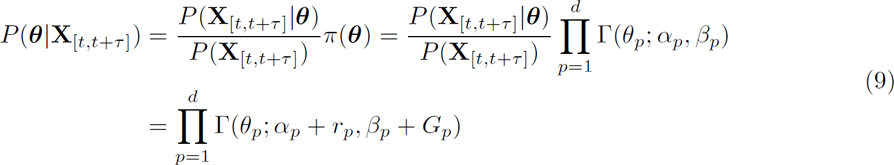

where *r_p_* is the number of reaction firings of reaction *p* on the interval [*t, t* + *τ*]. The term 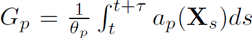 is the integral of the reaction propensity *a_p_* of the *p*^th^ reaction (Table A1), rescaled by the reaction constant *θ_p_*. The rescaling renders *G_p_* dependent only on the instantaneous configuration of the system at all points along the trajectory, and not on the reaction constants. Hence, a new sample for *θ* given the newly simulated trajectory (line 13 of Algorithm 1) can be generated by simply sampling from the updated gamma posterior (9); furthermore, the summary statistics *r_p_* and *G_p_* are sufficient for describing the posterior distribution of *θ_p_*. Thus, the full trajectories do not need to be stored, but instead the new simulations can be used to merely update the parameters of the posterior distribution of *θ* (i.e. set *α_p_* ← *α_p_* + *r_p_, β_p_* ← *β_p_* + *G_p_*), reducing storage requirements.

#### 2.3 STILT: Stochastic Inference on Lineage Trees

The particle filtering strategy described in Algorithm 1 is suitable for inference of the latent history of a single cell. However, if the cellular lineage is known it is possible to exploit the tree structure to improve the performance of the inference algorithm, for instance by constraining the range of possible initial values for daughter cells at the moment of division according to the state of the mother cell. This is more informative than assuming arbitrary distributions for the initial conditions of latent species, as required in Algorithm 1. Moreover, when incorporating the tree structure, the inferred parameter values are required to generate trajectories that have a high likelihood for multiple cells simultaneously as the cells proliferate. Such an approach is more e cient than performing inference on all cells simultaneously while neglecting the chronological and genealogical order owing to the less constrained initial conditions and lack of convergence of the parameter distributions.

STILT performs tree-based inference as outlined in Algorithm 2. We consider a tree comprised of *N_c_* total cells, with indices *j* = 1,…, *N_c_*. For the *j*^th^ cell there is a series of *N_j_* (possibly multivariate) measurements obtained at times 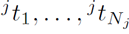, corresponding to each of the *N_o_* observed species, denoted by 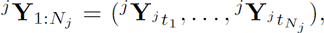. The algorithm begins by initializing a set of *K* particles for the tree’s founding cell with index *j* = 1, where each particle *k* comprises both a latent initial state 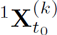 and a set of parameters *θ*^(*k*)^. All particles are initially equally weighted as 1/*K*. The algorithm then iterates through all of the measurement time points, where for simplicity we assume that measurements of all cells are obtained at a regular interval Δ*t* (i.e.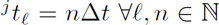); however, the method is equally valid for irregular measurement time intervals. At each iteration *i*, particles are resampled with frequencies proportional to their weights, and cells that are alive/measured at the current timepoint *i*Δ*t* are simulated one time step using the generative stochastic process with the sampled parameters *θ*^(*k*)^ to generate a sample 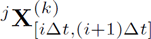 of the latent history of cell *j* over the time interval [*i*Δ*t,* (*i* + 1) Δ*t*].

If a cell is observed to divide between this timepoint and the next, the cell’s contents are allocated to the daughter cells. If both daughter cells are present, the total cellular contents of the mother cell must be conserved. For simplicity the division is assumed to take place just before the first observation of the two daughter cells at time (*i* + 1)Δ*t*. Cells are ordered such that cell *j* gives rise to cells with indices 2*j* and 2*j* + 1, with latent states 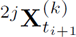 and 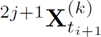, for the *k*^th^ particle. The conservation relationship between mother and daughter cells is enforced by requiring that 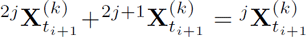. However, some species may be presumed to be identical between mother and daughter cells, e.g. DNA in active or inactive conformation.

After each forward simulation or division step the likelihood 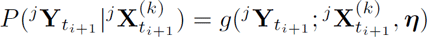 of each latent history is computed according to the observation function *g*, and used to reweight the particles. The observations of each cell are presumed independent conditional on the latent state, thus the likelihood of the complete set of observed cells is the product of the likelihoods of each cell, and the total weight of particle *k* is given by the product of the weights of each observed cell at that time point.

Assuming conditionally independent gamma priors *π*(*θ_p_*) = Γ(*θ_p_*; *α_p_*, *β_p_*) for each parameter *θ_p_*, the posterior probability of the model parameters conditional on the sampled latent histories until the current time point is shifted similarly to in (9), where *α_p_* increases by the summed number of reaction firings and *β_p_* by the summed integrals of the (rescaled) propensity functions, over *all* newly simulated trajectories on the interval [*i*Δ*t,* (*i* + 1) Δ*t*]. We define the set *A_i_* to be the set of indices of all cells observed at any point on the interval [0, *i*Δ*t*]:

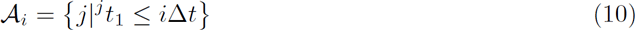

Let *^j^r_p_*(*i*Δ*t*) be the number of firings of reaction *p* in cell *j* at all times *t* ≤ *i*Δ*t* and 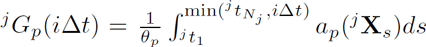 be the integral of the rescaled propensity function of reaction *p* for cell *j* until time *i*Δ*t*, for a particular realization of the stochastic process for cell *j*. With these definitions, the posterior joint density of model parameters *θ* is given by:

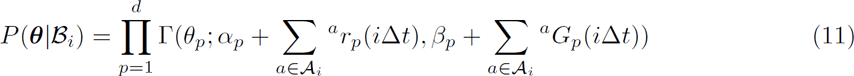

where the set 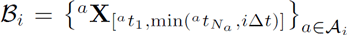 gives the set of realizations of the stochastic process for all cells observed at or before time *i*Δ*t*. Eq. (11) provides the means to generate samples *θ*^(*k*)^ from the probability density of model parameters conditional on a particular sampled complete genealogy 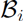. The parameter samples *θ*^(*k*)^ for each particle *k* are obtained by substituting the sampled trajectories for that particle into all expressions, i.e. *^a^***X** becomes *^a^***X**^(*k*)^, *^a^r_p_* becomes 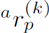, and *^a^G_p_* becomes 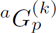. Since only the summary statistics are necessary to compute the posterior of the parameters, the full trajectories do not need to be saved, leading to a significant reduction in storage requirements. Finally, after iterating through all timepoints, the particles are resampled according to their weights yielding a set of *K* latent trajectories (if stored) and parameter sets. Thus the tree-based inference algorithm extends the single-cell-based inference algorithm (Algorithm 1) by establishing continuity between mother and daughter cells and initializing new latent trajectories for daughter cells according to the division process.

##### Algorithm 2: STILT algorithm

**Data:** A set of observations 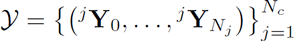 of *N_c_* cells observed at timepoints 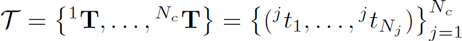; parameter prior *π*: ℝ*^d^ →* ℝ_0,+_; observation function *g*(**Y**; **X**, ***η***) = *P* (**Y***|***X**); number of particles *K*; measurement time interval Δ*t*

**Result:** A set of particles 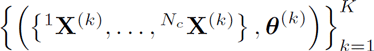 sampled from the posterior density 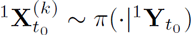

initialization;
**for** *k=1… K* **do**

Sample parameter values from the prior: *θ*^(*k*)^ ~ *π*(*θ*);
Sample initial state conditional on first observed data point: 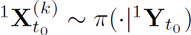;
Initialize particle weight to 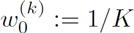
Generate a set of particle indices *ε*^(*k*)^ ∈ {1,…, *K*}, *k* = 1,…, *K* such that
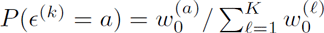;
compute maximum of all timepoints: 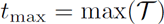;
loop over all observed timepoints;
**for** *i* = 0: [*t*_max_/Δ*t*] **do**

determine cells alive at this timepoint;
*σ* = {*j*|*i*Δ*t* ∈ *^j^***T**};
loop over particles;
**for** *k = 1… K* **do**

loop over cells at current timepoint;
**for** *j* ∈ *σ* **do**

Get index of current timepoint for cell *j*;
*ℓ* = find(*^j^t_ℓ_* = *i*Δ*t*);
Compute the partial weight of particle *k* for the *j^th^* cell: 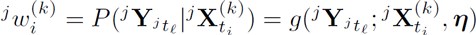;
Generate a sample trajectory 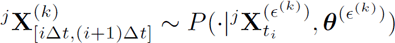;
**if** (*i* +1) Δ*t* ∉ *^j^***T then**

Initialize daughter cells;
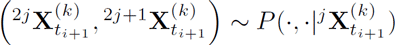;
**else**

Concatenate to previously sampled trajectory: 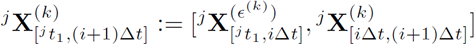;
Compute the total weights for particle *k*: 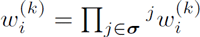;
Generate a set of particle indices *ε*^(*k*)^ ∈ {1,…, *K*}, *k* = 1,…, *K* such that
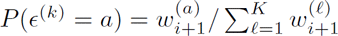;
Sample new parameters *θ*^(*k*)^ from the conditional density: 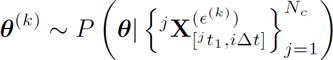;
Construct a sample of *K* particles from the posterior: 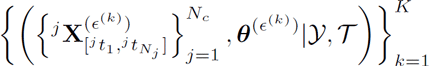

#### 2.4 Single cell versus tree-based inference

In the previous sections we presented two algorithms for inferring model parameters using the bootstrap particle filter. Algorithm 1 treats each cell individually while Algorithm 2 explicitly incorporates the known cellular lineage tree. As an alternative to Algorithm 2, it is also possible to fit all cells by simply discarding lineage knowledge, artificially synchronizing all cells to start at the same time point, and fitting cells in parallel. However, testing revealed that this approach quickly converges to local optima due to the inability to fit all cells simultaneously without information about their initial conditions. In contrast, Algorithm 2 benefits from exploiting the initial iterations of the algorithm with fewer cells in order to pre-converge the parameter distributions, and provides good estimates for the initial conditions of daughter cells upon division of the mother cell under the assumed division process.

The single-cell based inference performed consistently worse than the tree-based inference using STILT on synthetic data (Table S3, Figure S4), in terms of model identification and parameter estimates. This is likely because the single cell-based inference does not exploit the lineage structure to improve the estimation of the initial conditions (i.e. by enforcing conservation of inherited cellular material between mother and daughter cells), and because it is not obvious how to combine the inference results of individual cells in order to provide a better estimate of the overall population parameters.

### 3 Model assumptions

#### 3.1 Cell division

We assume that at the time of division, each mother cell allocates its contents (mRNA and protein) randomly with equal probability to each daughter cell. Thus the number of mRNA molecules are distributed as:

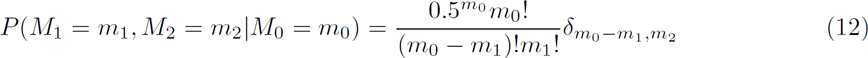

where *M*_1_, *M*_2_ are the number of mRNA molecules of the two daughter cells upon division, and *M*_0_ that of the mother cell; *δ_x,y_* is the Kronecker delta. The protein contents are allocated analogously, although for numerical reasons the binomial distribution is approximated by a normal distribution with equivalent mean and variance.

The conformation of the DNA (i.e. active or inactive) for each gene is assumed to persist from mother cell to daughter cell at division. This assumption is motivated by the observation that progeny of a cell typically resemble the ancestor cell in terms of gene activity. However, due to the stochastic nature of the model, some simulated trajectories may still switch activation state shortly after division, e ectively permitting cells to also switch activation state upon division if this trajectory exhibits high likelihood. We note however that this is not an essential assumption of the inference procedure and can be easily changed for alternative scenarios.

#### 3.2 Feedback models

In the feedback models, the DNA activation and inactivation rates are modified by the protein abundance. For the Negative Feedback model the amount of protein modulates the rate of DNA inactivation and for the Positive Feedback model protein modulates the rate of DNA activation. We consider the case of switch-like activation/inactivation of the DNA with increasing concentration of protein. We achieve this by assuming that the propensity of DNA activation in the Positive Feedback model is given by *a*_on_ = *k*_on_*P* ^2^ and of DNA inactivation by *a*_off_ = *k*_off_ *P*^2^ in the Negative Feedback model.

We assume fast binding and dissociation of protein to the DNA relative to protein production and degradation, such that the protein abundance can be treated as approximately constant on the time scale of binding dynamics. With this assumption, the probability of DNA being in either the active (DNA*) or inactive state (DNA) evolves for the Negative Feedback model according to the chemical master equation:

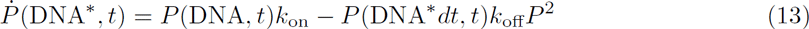

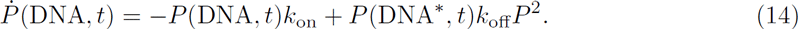

Requiring that *P*_SS_(DNA*) + *P*_SS_(DNA) = 1, the steady state solution gives:

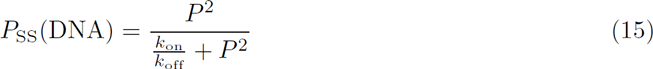

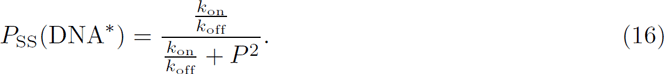

Thus the probability of DNA being inactive is a sigmoidally increasing function of the number of proteins. This activation function is a Hill function with coefficient 2, corresponding to cooperative binding of two Nanog molecules at the promoter/enhancer. The quantity (*k*_off_/*k*_on_)^1/2^ determines the protein abundance for which the DNA has 50% probability of being active.

Similarly, for the Positive Feedback model, the probability of the DNA states evolves as:

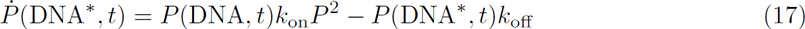

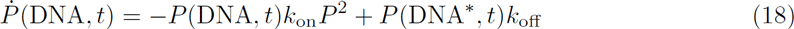

for which the steady state solution gives:

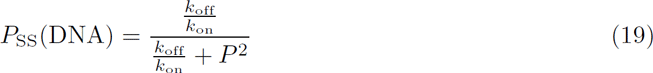

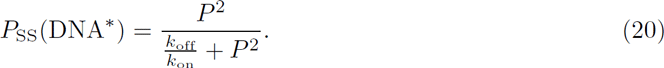

which is a Hill function with coefficient 2 for the probability of DNA activation.

#### 3.3 Biallelic expression

In the synthetic data sets we consider expression dynamics of a single allele only. Thus, there are precisely four species: DNA in active and inactive conformations, mRNA and protein. In the NanogVENUS subtree modeling, the fluorescent fusion protein NanogVENUS is also expressed only in a single allele [8]. Hence we apply the same models as for the synthetic data. However, there is also the possibility of expression in the other, unlabeled Nanog allele. Since we cannot quantify this allele, we assume that its expression is highly correlated to the observed allele, which has previously been reported for the same system [9]. Assuming equal proportions of observed and unobserved Nanog protein, the total amount is roughly double, which translates to a fourfold rescaling of the estimated rate constants *k*_on_ and *k*_off_ for the Positive and Negative Feedback models, respectively. The inference procedure is otherwise not affected.

#### 3.4 Cellular compartments

For simplification, we do not explicitly model cellular compartments such as cytoplasm or nucleus. Thus nuclear translocation effects are implicitly captured by the estimated rate constants. The NanogVENUS experiments analyzed [8] quantify only nuclear protein. Thus the model pertains only to the expression dynamics of nuclear Nanog. The estimated protein degradation rate also captures both degradation and implicitly the dilution to the cytoplasmic compartment.

### 4 Model specifications

#### 4.1 Measurement function

STILT requires the specification of an observation function that yields the likelihood of a particular observation 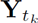 given the state of the latent history 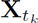 at some time *t_k_*, this function is referred to as *g* in Section 2 and corresponds to the measurement process. In the case of time-lapse fluorescence microscopy one typically assumes that the fluorescence intensity is proportional to the abundance of fluorophores. Assuming that the measurement process induces some small error *ε*, this gives the simple linear relation

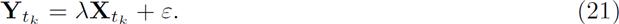

where *λ* is the mean fluorescence intensity per molecule. In the analysis of the synthetic lineage trees (Figure 2), no conversion between proteins and fluorescence intensity was necessary, i.e. *λ* = 1. We let 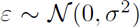 with standard deviation *σ* = 200 proteins. This is the same as was used for generating the noisy observations of the synthetic data.

In the analysis of NanogVENUS fluorescence lineage trees, we estimate *λ ≈* 10^5^ proteins per unit fluorescence intensity based on comparison of mean concentration estimated from Western blot and mean fluorescence intensity of unbiased NanogVENUS lineage trees (see Section 7.2). We likewise assume Gaussian measurement noise, and use the NanogVENUS lineage trees to estimate the standard deviation to be approximately *σ* = 2 × 10^4^ based on the small signal fluctuations. We note that the exact value of the measurement error should not bias the inference results, but rather, too small a value will lead to non-robust estimation of parameters as there is a higher risk of too many particles being discarded due to low likelihood, and too high a value leads to a poorer ability to infer model parameters as too few particles are discarded. However, the robust estimation of model parameters and apparent divergence from the prior (see Figure S8) seems to indicate an adequate choice for the magnitude of the measurement error *σ*.

#### 4.2 Prior distributions

STILT is a Bayesian inference technique and thus requires specification of prior distributions for model parameters. We utilize Gamma priors distributions for each parameter, which greatly simplifies the sampling procedure (see 2.2). For the *in silico* experiments, the true model parameter were known. In this case, the priors distributions were chosen so that they i) contain the true model parameters, and ii) allow for easy visual assessment of convergence to the true model parameters.

For the investigation of the Nanog subtrees, the prior distributions were obtained as follows.

##### mRNA degradation

The half-life of Nanog mRNA in mouse ES cells cultured in serum/LIF has been estimated as 6.8 h [10], and 3.9-6.4 h [11]. We thus chose *α* = 8 and *β* = 40 for which the 95% confidence interval of the half-life is (1.9 h, 8.0 h).

##### Transcription

The Nanog mRNA transcription rate was recently estimated as approximately 125 molecules/h in serum/LIF [10]. Moreover, the number of Nanog mRNA molecules in mouse ES cells under serum/LIF conditions is approximately 300 or fewer, rarely exceeding 400 [12, 13]. Using the mean estimated degradation rate of 0.2 h^−1^, and assuming DNA remains active, the expected number of mRNAs (given by *k_m_/g_m_*) would thus be approximately 625 molecules which is more than typically expected. We therefore set the prior distribution constants to be *α* = 10, *β* = 0.1 for which the 95% confidence interval of the transcription rate becomes [47.95, 170.85] h^−1^, and the expected number of transcripts is approximately 500. This number is somewhat reduced by the fact that the DNA is typically not persistently active, and by the cell division process which reduces the mRNA count by a factor of approximately 2.

##### Protein degradation

The half-life of Nanog has been reported as approximately 2.1 ± 0.8 h [14]. Nanog’s half-life was also measured in the analyzed data set and found to be closer to 5 h [8]. We therefore conservatively set the prior distribution constants to be *α* = 2, *β* = 4 such that the 95% confidence interval of the half-life is approximately 0.5 *−* 11.5 h.

##### Translation

The translation rate of Nanog is not well characterized. However, the estimated mean number of Nanog molecules per cell is approximately 350,000, the mean degradation rate approximately 0.2 h^−1^, and mean number of mRNA molecules roughly 200. In the deterministic limit, the expected number of proteins is given by 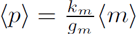, where 〈*p*〉 and 〈*m*〉 denote the mean protein and mRNA counts, respectively. Substituting our estimates, we obtain approximately 350 mRNA^−1^ h^−1^. We thus chose *α* = 3, *β* = 0.01 such that the mean (std.) translation rate is 300*±*173 mRNA^−1^ h^−1^. We note that this is in rough agreement with the mean estimated translation rate of 478 mRNA^−1^ h^−1^ over all analyzed genes in mouse fibroblasts (maximum estimated rate of 1000 mRNA^−1^ h^−1^) [15].

##### DNA activation and inactivation

The rate of DNA activation and inactivation for Nanog is not well studied. By inspecting the analyzed NanogVENUS subtrees, we surmise that periods of rapid fluorescence intensity likely correspond to periods of DNA activity, and periods of decline to DNA inactivity. Thus activation and inactivation presumably proceeds with expected waiting time on the order of hours and not days or longer. Consistent with this, Sokolik *et al.* estimated active/inactive switching times to be approximately 3.8 *±* 1.2 h [14]. Since the autoregulation models studied di er in the form of their activation/inactivation rates, it was necessary to choose priors for each separately. For each of the three models, the prior distributions for *k*_on_ and *k*_off_ were chosen such that the waiting times for activation/inactivation were on the order hours, and such that trees simulated with these parameters produced reasonable dynamics, i.e. observed DNA state switching, and approximately correct order of magnitude for number of proteins.

#### 4.3 Initial conditions

The initial state of the founder cell of the cellular lineage trees is unknown. In the considered models DNA and mRNA are entirely latent, while protein is observed with noise. For both the *in silico* experiments and the NanogVENUS subtrees, we initialized each cell’s DNA state to be active or inactive with 50% probability, and to have an initial mRNA count uniformly sampled from [0, 50]. The protein copy number was sampled from a Gaussian distribution centered on the first observation, with standard deviation specified by the measurement function (see Section 4.1).

#### 4.4 Number of particles

All implementations of the bootstrap particle filter require specification of the number of particles to use for approximating the latent history and posterior parameter distributions. The accuracy of the approximation improves in the Monte Carlo sense as the number of particles is increased. However, the incurred computational overhead increases proportionally. We used 7 × 10^5^ particles for inference of the synthetic lineage trees, 10^5^ for each synthetic cell for the single-cell-based algorithm, and 10^6^ particles for the NanogVENUS subtrees. We determined the number of particles to use based on robustness of convergence of the parameter posteriors, and run-time. Inference typically completed on a multicore machine in approximately 10h for 10^6^ particles for a single subtree.

### 5 Implementation

STILT (see Algorithm 2) was implemented using Matlab 2015a. It includes code for importing SBML models and fast, parallel stochastic forward simulations for the system state using Matlab’s parallel computing toolbox.

#### 5.1 Model definition via SBML

Our implementation supports the import of biochemical network models from SBML using libS-BML 5.12.0 [16], but can also be specified directly in Matlab. Species, reactions and their parameters are translated into a stoichiometric matrix and vectorized Matlab functions for computing reaction propensities (Section 1).

#### 5.2 Simulation

The stochastic simulation code was implemented using explicit, adaptive *τ*-leaping [17, 18], which generates approximate samples from the exact stochastic process. In general, *τ*-leaping approximates the Markov jump process by a Poisson process with the same expected number of reactions firing for time intervals where the reaction propensities remain relatively constant. Such an approximation is generally necessary when the system becomes stiff, i.e., when there exist reactions with widely varying time scales such as is the case for the protein production and degradation compared to DNA activation and inactivation. As for ref. [17] our implementation distinguishes between critical and non-critical reactions (based on current educt availability, the parameter *N_c_* is set to 10) and performs explicit *τ*-leaping (with parameter *ε* = 0.03) for non-critical reactions with error bounding implemented for first, second and third order reactions. The simulation code was implemented completely vectorized and provides the approximated integrated reaction propensities (for mass action propensities only) and the number of reactions firing, which are required for inference (see Section 2). The forward simulation code can be further accelerated by converting the entire function or only the Poisson random number generation to C code, which lead to significant speedup for the studied systems.

#### 5.3 Data structure

Measurement data are specified using a generic Matlab structure containing measurement times, cell number and an indicator for censoring (e.g. for inaccurate or missing data) as well as measurements and their respective measurement errors. Field names of measurements are automatically matched to SBML species. Parameter priors, model specifications (e.g. behaviour on cell division), compilation behavior and other user configurations are provided via an options structure.

### 6 Model evaluation

#### 6.1 Marginal likelihoods and Bayes Factors

The particle filtering approach presented above can be used for performing model comparison via Bayes Factors, i.e. by computing 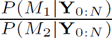, the ratio of the posterior probabilities of Model 1 (*M*_1_) to Model 2 (*M*_2_) for any two models. As before, we denote the series of observations at times *t*_0_,…, *t_N_* by 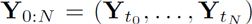. Using Bayes’ law, one can reformulate the marginal posterior probability of a model M as:

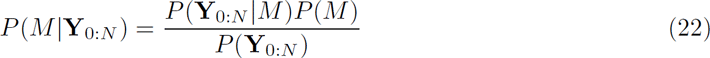

Following Wilkinson *et al.* [2], p. 294, we can approximate the marginal likelihood of the model *P* (Y_0:*N*_|*M*) using the sampled particles at each iteration *i*. Firstly, the distribution of the observed data at time *t_i_*_+1_ depends only observations up to *t_i_*: 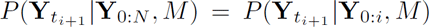. Moreover, this probability is approximated by the expectation of the likelihood, or weights *w*^(*k*)^, of the particles:

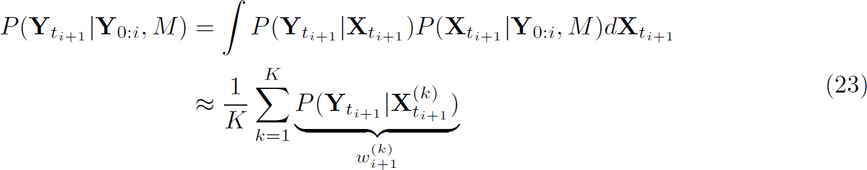

where the 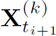 are sampled (via the particle filter) from the marginal posterior up to time *t_i_*_+1_ given by 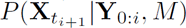. This is nothing more than a Monte Carlo approximation of the integral, which provides an unbiased approximation of 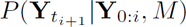 with variance decreasing as *K*^−1^ [19].

Next, since the distribution of each observation depends only on previous observations, the marginal probability of the entire set of observations *P* (Y_0:*N*_|*M*) is given by the product:

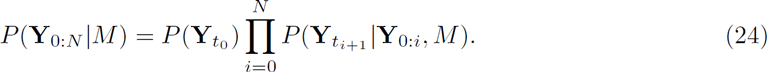

Assuming *a priori* equally likely models, the factor of *P* (*M*) in (22) cancels between the two models and the Bayes Factor reduces to the ratio of marginal likelihoods. In the analysis presented in this work we primarily utilize log Bayes Factors and marginal log likelihoods due to their superior numerical performance.

#### 6.2 Goodness-of-fit test

To assess the extent to which a particular model agrees with an observed data set, we developed a simple goodness-of-fit (GOF) test. The GOF test utilizes an estimate for model parameters obtained from the particle filter to generate many synthetic datasets, which are then compared against the measured data. Specifically, we use the assumed model to generate many synthetic lineage trees of the same number of generations as the observed data using the median posterior parameter estimate of each parameter. For each newly simulated data set, we approximate its log likelihood conditioned on the parameter set that was used to generate that data. The conditional log likelihood (CLL) is approximated again via a particle filter, where the parameters are fixed. This conditional particle filter only samples from the latent history of all state variables while keeping the parameters fixed, and is essentially the same as in Algorithm 2 omitting the parameter resampling step.

To compensate for the fact that the simulated datasets and the measured dataset do not necessarily contain the same number of transitions, we normalize the estimated CLL of each simulation by the number of simulated transitions (i.e. between measurement time points). We likewise normalize the CLL of the actual data by the number of transitions (subtracting censored observations), to obtain the average CLL per transition. Without this compensation the CLL is always decreasing with the number of transitions since the log likelihood is never greater than zero, which could potentially bias the CLL depending on the random lifetime of each simulated cell.

In all applications of the GOF we used 300 samples to approximate the null distribution of the CLL, and 500 particles per sample to approximate the CLL. The GOF test approximates the null distribution of the CLL, i.e. the distribution of CLL values yielded by the particle filter when the parameters and model utilized are known to be true. We compute the CLL of the actual dataset using the same parameter values and compare it with the null distribution of the CLL. If it lies within this distribution, then with high probability the dataset could have been produced by the chosen mechanistic model and parameter set, and the model cannot be rejected. Conversely, if the CLL of the observed dataset lies outside the null distribution, the model and parameters are unlikely to have produced this dataset. Thus, we define three categories of model agreement with the null distribution: reject (*p* < 0.01 or *p* > 0.99), marginal (*p* < 0.025 or *p* > 0.975), and accept, otherwise. Empirical *p*-values are estimated using the empirical cumulative distribution function of the estimated CLL.

### 7 Time-lapse fluorescence microscopy data

#### 7.1 Pre-processing

We obtained quantified time-lapse fluorescence microscopy movies of NanogVENUS in mouse embryonic stem cells from the data set of Filipczyk *et al.* [8] and converted fluorescence intensities to protein numbers as described in Section 4.1.

Since the time-lapse fluorescence microscopy quantification introduces error due to variability in the cellular (nuclear) segmentations, background correction, etc., we performed a data cleaning step prior to analysis. We censored measurements for all automatically segmented cells that could not be manually verified, e.g. if the cells were too densely packed or overlapping to be reliably quantified. We further censored very large jumps (the top 5% of absolute change in intensity) in the quantified intensity of individual cells, which result from either contamination due to microscopic debris, errors in cell segmentation, and in some cases jumps in the intensity at the last time point before cell division which presumably arise due to a sudden change in the cellular morphology preceding division that leads to a large overlap of cytoplasmic and nuclear volumes (see Supplementary Fig. 5 for examples). Censored measurements a ect the inference by rendering all simulated particles equally likely at that iteration; the algorithm otherwise proceeds as normal.

After cleaning and selecting data sets, we obtained a total of 7 quantified cellular genealogies from 3 different experiments. To improve computational e ciency, and for comparison with the synthetic data, we subdivided these large trees into smaller subtrees each containing 7 cells with no overlap between subtrees, thus obtaining 4 subtrees from the first experiment, 8 subtrees from 3 di erent parent trees of the second experiment, and 1 subtree from each of 3 parent trees of the third experiment; in total 15 subtrees were used for further analysis (see Supplementary Fig. 7).

#### 7.2 Estimation of fluorescence intensity conversion factor

To estimate the absolute number of NanogVENUS molecules per cell, we performed Western Blots experiments on 10% polyacrylamide gels. We compared Western Blots with a known quantity of NanogGFP single knock-in fusion ESCs and with di erent quantities of recombinant GFP (Catalogue number: 632373, Clonetech, CA, USA). Both NanogGFP and GFP proteins were detected using an anti-GFP primary antibody consisting of two monoclonal clones (Catalogue number: 1181460001, Roche, Mannheim, Germany). Western Blot band intensities were quantified by using the Gel Analyser tool in FIJI to gate on protein lanes and quantify band intensities over background. We found that the relationship between the GFP quantities *x* and the corresponding intensity *y* is best described by a sigmoid function:

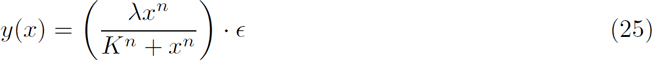

The model parameters *λ_j_*, *K_j_*, *n_j_* were obtained by local optimization using multiple restarts initialized according to Latin-hypercube sampling. The exponent *n* determines the shape of the sigmoid function, *K* sets the inflection point, *λ* is the maximum of the curve and *ε* is a log-normally distributed error term with expectation 1 and standard deviation *σ*, as is suggested for Western Blot data [20]. We compared this model against linear models both with and without intercept and found it to be superior according to both the Bayesian Information Criterion and coe cient of variation between replicates. We solve Eq. (25) for *x* to obtain

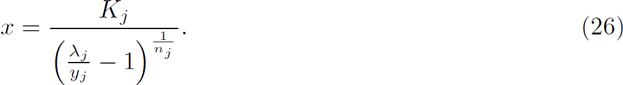

The total quantity of protein *x* is related to the cellular average *P_j_* as

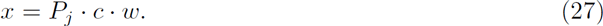

where *c* is the number of loaded cells and *w* is the molecular weight for the protein of interest. Thus we determined the number of proteins *P_j_* per cell from the sample intensity *y_j_* of each Western Blot replicate *j* by first computing *x* from the observed intensity *y* according to Eq. (25), and then substituting into Eq. (27).

As *P_j_* is a combination of uncertain variables, we obtained error bars for each *P_j_* individually by applying standard error propagation to account for uncertainties in the number of cells *c* (we assume a standard deviation of 10%) and uncertainties in the model (estimated via the standard deviation *σ* of our noise model). However, we find that the uncertainties for each individual replicate *P_j_* are always smaller than the inter-replicate standard deviations by a factor of 0.3 or smaller. Therefore, we only consider the standard deviation across replicates, as this is the dominant source of uncertainty in our procedure. Finally, we determined the fold-change between NanogVENUS and NanogGFP from three Western Blots. Uncertainty of protein abundance over replicates was estimated by simple error propagation. All above experiments and analysis were performed in triplicate.

The resulting estimate is of approximately 350, 000 *±* 72, 000 molecules of NanogVENUS expressed in each cell. Using the distribution of NanogVENUS fluorescence intensities over unsorted mESCs, we obtain a median intensity of approximately 3.5, from which we determine the calibration factor of NanogVENUS fluorescence intensity to NanogVENUS molecules count to be approximately 100,000.

### 8 Experimental validation of Negative Feedback model

#### 8.1 Exogenous Nanog construct

##### 8.1.1 ESC culture

Mouse ESCs were of R1 background and report endogenous Nanog protein levels from one allele by fluorescent fusion proteins, either NanogVENUS (NV) or NanogKATUSHKA (NK). ESCs were cultured in DMEM (Catalogue number: D1145 Sigma, MO, USA) supplemented with 2mM GlutaMAX (Catalogue number: 35050-038, Gibco, USA),1% Non-essential amino acids (Catalogue number: 11140-035, Gibco, CA, USA), 1mM Sodium Pyruvate (Catalogue number: S8636, Sigma, MO, USA), 50uM b-mercaptoethanol (Catalogue number: M6250, Sigma-Aldrich, USA), 10% FCS (Catalogue number: 2602P250915, PAN, Aidenbach, Germany) and 10ng/ml LIF (GFM200, Cell Guidance Systems, Cambridge, UK) on 0.1% porcine gelatin (Sigma, Catalogue number: G1890-100G).

##### 8.1.2 Nanog overexpression experiments

30000 cells were seeded in a well of a 24w plate and transfection was performed 5-7h after seeding. For transfections, 250ng plasmid, 1ul Lipofectamine 2000 (11668-019, Life Technologies) and 50ul Opti-MEM (31985-062; Life Technologies) were mixed and added to the cells. Cells were analyzed by flow cytometry 46h after transfection using a BD LSR Fortessa (BD Biosciences, CA, USA) and data were analysed with FlowJo (OR, USA). Cells were gated for non-debris and singlets using FCS-A, SSC-A and FCS-W channels. Fluorescence channels were compensated using controls that only expressed one of each fluorescent protein. R1 wild-type cells were used as control for cellular autofluorescence. All experiments have been performed as triplicates.

##### 8.1.3 Expression plasmids

The Nanog coding sequence was cloned in several variants as a 2A construct into a piggybac vector that has been modified to express a fluorescent nuclear membrane tag (iRFPnucmem) from the CAG promoter using the In-Fusion system (Catalogue number: 638911, Takara, Japan). The resulting constructs are supposed to express iRFPnucmem and Nanog proteins in equal abundances. The plasmid CAG.iRFPnucmem-P2A-NanogVENUS was used in overexpression experiments to allow for comparison of exogenous NanogVENUS levels with endogenous NanogVENUS levels of the R1 NanogVENUS cell line. The NanogVENUS plasmid performed identically to positive control plasmids (CAG.iRFPnucmem-P2A-Nanog; with or without ATG for Nanog). An empty vector control (CAG.iRFPnucmem-P2A) was also used during experiments.

#### 8.2 Exogenous NanogVENUS expression compartments

Using the wild-type cell line which expresses no NanogVENUS, we obtained the NanogVENUS intensity distribution due only to autofluorescence (Figure S10A), from which we deduce the 0.95 quantile of NanogVENUS autofluorescence, denoted 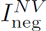. We then used the expression distribution of a cell line which expresses NanogVENUS at one endogenous allele (NV cell line, see Section 8.1) to derive the 0.95 quantile of unperturbed NanogVENUS expression, denoted 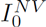 (Figure S10A).

We define the NanogVENUS No Exogenous compartment as NanogVENUS fluorescence intensities that are below 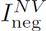. Since NanogVENUS is only expressed on one allele of the NV cell line, the total quantity of Nanog in the cell is approximately twice this amount. Based on this we define the 1x overexpression (OE) compartment to be intensities that are above the No Exogenous compartment but below 200% of the normal level 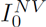. The 2x OE compartment has NanogVENUS intensities *I^NV^* between 2-4 times normal, and the 3x OE compartment between 4-6 times normal. Cells with higher NanogVENUS intensities fall into the “very high” intensity compartment.

#### 8.3 Downregulation of endogenous Nanog levels

To investigate negative feedback, we utilize the NK cell line (see Section 8.1), and compute the expression levels of endogenous NanogKATUSHKA for di erent levels of exogenous transgenic NanogVENUS expression. We first obtain the autofluorescence intensity distribution on the NanogKATUSHKA channel using NV cells which express no KATUSHKA, from which we estimate the 0.95 quantile of intensity of the KATUSHKA negative compartment, denoted 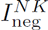 (Figure S10B). We first normalize NanogKATUSHKA fluorescence relative to background by subtracting 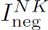 from the measured intensities. We then compute the median fold-change, for each overexpression compartment *k*, of normalized expression of NanogKATUSHKA relative to that of the No Exogenous compartment (see Figure 4E, Table A2). Denoting the NanogKATUSHKA expression of a cell with index *j* by 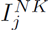, and its NanogVENUS expression by 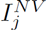, the median fold-change for compartment *k* is given by:

**Table A2:**
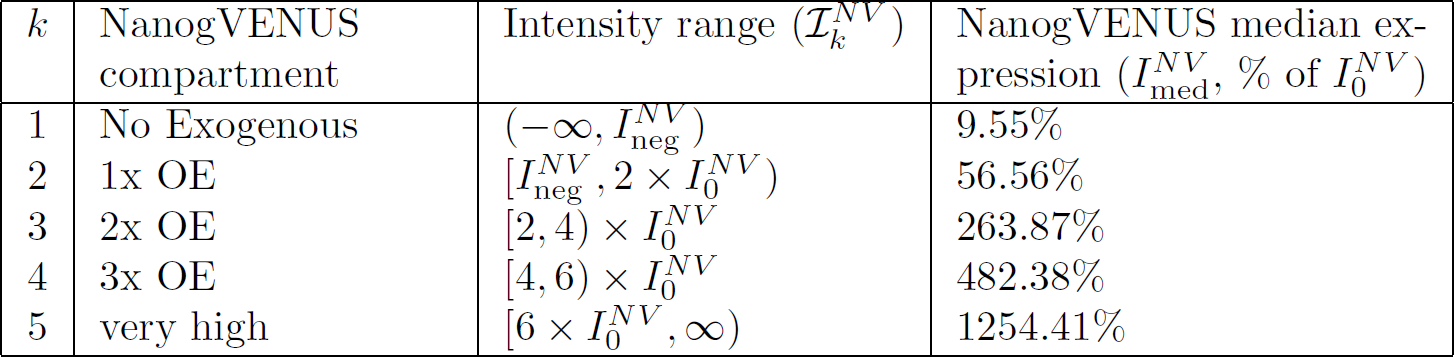
NanogVENUS compartments. Intensities are defined relative to the 0.95 quantile of NanogVENUS expression in the NV cell line 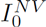, and the 0.95 quantile of NanogVENUS expression in the WT line, 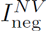. For each NanogVENUS intensity compartment we compute the median NanogVENUS intensity 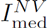 relative to 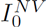.

**Table A3:** Parameter values used for simulation of exogenous Nanog.

| Parameter | Value |
| --- | --- |
| $k_{\text{on}}$ | 0.6854 |
| $k_{\text{off}}$ | $5.690 \times 10^{-11}$ |
| $k_m$ | 110.3 |
| $g_m$ | 0.226 |
| $k_p$ | 619.5 |
| $g_p$ | 0.220 |

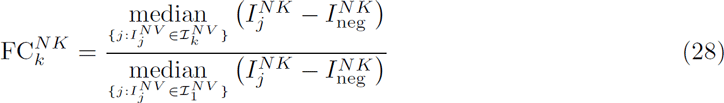

##### 8.3.1 Comparison of experimental replicates

The NanogVENUS overexpression experiment described above was performed three times. To compensate for batch effects, the distributions in each experiment were normalized relative to the first experiment. Specifically, for each replicate, the fluorescence intensity of endogenous NanogKATUSHKA was scaled linearly so that the median intensity of cells matches to the median intensity of the first experiment. The same NanogKATUSHKA background level 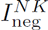 was used for each of the three replicates (see Section 8.3).

#### 8.4 *In silico* perturbation experiment

We replicate the experimental setup by extending the Negative Feedback model to include exogenous Nanog (*P*_ex_), such that the propensity of DNA inactivation becomes

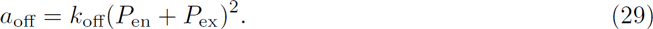

From the time-lapse fluorescence microscopy movies of NanogVENUS subtrees we obtain estimates of posterior distributions of parameters for the Negative Feedback model (Table S5b). We compute the median of the posterior for each parameter and subtree, and then the mean of the medians for each parameter over the subtrees (Table A3). We then perform *in silico* perturbation experiments using these mean parameter values and various levels of exogenous Nanog.

To mimic the experimental setup, we sampled intensity values 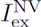 directly from the measured distributions of exogenous NanogVENUS for each overexpression compartment separately. We convert the sampled intensities into a specific number of molecules by computing the overexpression relative to wild-type NV cells. Since the fluorescent reporter is expressed only on one allele, a 100% increase of NanogVENUS corresponds to an approximately 50% increase in total Nanog levels. We assume approximately 2 × 10^5^ NanogVENUS molecules in a cell with no exogenous perturbation (see Figure S7). We thus compute the sampled amount of exogenous NanogVENUS molecules as 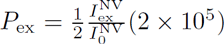.

For each sampled quantity of exogenous NanogVENUS, we generated 50 synthetic lineage trees of 5 generations each. The founder cell of each lineage tree had DNA initially active, between 0 and 150 mRNA molecules (uniformly sampled), and between 10^5^ and 2 × 10^5^ Nanog molecules (uniformly sampled). The exogenous Nanog levels were held fixed at the sampled value for the duration of the simulation.

#### 8.5 Comparison to simulations

The distribution of endogenous Nanog following perturbation was computed for each overexpression compartment after 46h of simulated time. The fold-change of endogenous Nanog expression relative to expression with no perturbation was computed analogously to (28):

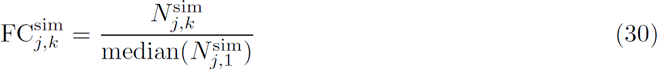

where 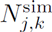 denotes the number of endogenous Nanog molecules of cell *j*, 46 hours after the perturbation corresponding to compartment *k*.

In Figure 4E, we plot the distribution of the fold-change of simulated cells with respect to the No Exogenous compartment as a box-and-whiskers plot (median shown as red line). We compare this against the median (mean, s.e.m., n=3 experimental replicates) fold-change computed for the experimental data, see Eq. (28). The comparisons against each experimental replicate individually are shown in Figure S10E.

